# Excitatory and inhibitory effects of HCN channel modulation on excitability of layer V pyramidal cells

**DOI:** 10.1101/2022.03.30.486368

**Authors:** Tuomo Mäki-Marttunen, Verónica Mäki-Marttunen

## Abstract

Dendrites of cortical pyramidal cells are densely populated by hyperpolarization-activated cyclic nucleotide-gated (HCN) channels, a.k.a. *I_h_* channels. *I_h_* channels are targeted by multiple neuromodulatory pathways, and thus are one of the key ion-channel populations regulating the pyramidal cell activity. Previous observations and theories attribute opposing effects of the *I_h_* channels on neuronal excitability due to their mildly hyperpolarized reversal potential. These effects are difficult to measure experimentally due to the fine spatiotemporal landscape of the *I_h_* activity in the dendrites, but computational models provide an efficient tool for studying this question in a reduced but generalizable setting. In this work, we build upon existing biophysically detailed models of thick-tufted layer V pyramidal cells and model the effects of over- and under-expression of *I_h_* channels as well as their neuromodulation by dopamine (gain of *I_h_* function) and acetylcholine (loss of *I_h_* function). We show that *I_h_* channels facilitate the action potentials of layer V pyramidal cells in response to proximal dendritic stimulus while they hinder the action potentials in response to distal dendritic stimulus at the apical dendrite. We also show that the inhibitory action of the *I_h_* channels in layer V pyramidal cells is due to the interactions between *I_h_* channels and a hot zone of low voltage-activated Ca^2+^ channels at the apical dendrite. Our simulations suggest that a combination of *I_h_*-enhancing neuromodulation at the proximal apical dendrite and *I_h_*-inhibiting modulation at the distal apical dendrite can increase the layer V pyramidal excitability more than any of the two neuromodulators alone. Our analyses uncover the effects of *I_h_*-channel neuromodulation of layer V pyramidal cells at a single-cell level and shed light on how these neurons integrate information and enable higher-order functions of the brain.

## 1 Introduction

In the brain, higher-order cognition and consciousness are believed to rely on the highly specialized neurons that populate the cortex, the layer V pyramidal cells (L5PCs). Thanks to their complex morphology and connectivity, feed-forward sensory-related stimuli and feed-back context-dependent inputs arrive at spatially distinct sections of the L5PC dendritic tree [Larkum, 2013]. The different inputs are integrated in the soma and together determine the specific patterns of activity of the neuron. The effect that feedback context-dependent inputs have on somatic excitability is partly mediated by hyperpolarization-activated cyclic nucleotide-gated (HCN) channels, a.k.a. *I_h_* channels. L5PCs strongly express *I_h_* channels in their apical dendrite that reaches up to layer I of the cortex [Berger et al., 2001]. Moreover, the apical dendrite of L5PCs is abundantly projected to by neuromodulatory terminals from subcortical regions, including VTA (dopaminergic NM) [Gulledge and Jaffe, 1998], basal forebrain (cholinergic modulation) [Turrini et al., 2001], and locus coeruleus (noradrenergic modulation) [Branchereau et al., 1996]. The interplay between *I_h_*-driven communication between apical and somatic sections, together with neuromodulatory input, confers L5PC neurons their complex integrative capacity, but the details of the mechanisms behind this interplay remain an open question.

*I_h_* channels give the neuron an extensive set of modes of excitability. A reason for this is that they can either depolarize and hyperpolarize the cell membrane during subthreshold membrane potential fluctuation. That is, their reversal potential lies between −45 and −30 mV [Hestrin, 1987, McCormick and Pape, 1990] and therefore their effect on membrane excitability will depend on the electrical environment. In addition, the *I_h_* channels can be modulated by several neuromodulators such as dopamine, acetylcholine and norepinepherine [Ballo et al., 2010, O’Donnell et al., 2012, He et al., 2014]. Previous experimental work has assessed separately the effects of *I_h_* blockage or neuromodulation in terms of their effect on somatic [Pedarzani and Storm, 1995] and dendritic [Tsay et al., 2007, Labarrera et al., 2018], excitability. However, the exact way in which the concurrence of the multiple factors affect the direction of *I_h_* modulation on neuron excitability remains unclear. Computational modelling offers the possibility to assess the mechanisms behind cellular electrical properties, and generate testable predictions. In this work, we used biophysically detailed computational modelling to analyze the effect of *I_h_* channels and their neuromodulation on L5PC activity.

To analyze the effects of *I_h_* channels and the way they modulate L5PC excitability, we used existing biophysically detailed neuron models of thick-tufted L5PCs with reconstructed morphologies. We determined the threshold currents or conductances for many types of stimulus protocols in presence and absence of *I_h_* currents. In this way, we characterized the types and locations of stimuli for which *I_h_* currents are excitable and those for which they are shunting (i.e., inhibitory). We also modelled the response of the neuron when the *I_h_* channels were modulated by dopamine or acetylcholine by introducing experimentally observed effects of these neuromodulators on the voltage-dependence profile of *I_h_*. We showed that *I_h_* channels shunt stimulation at the distal apical dendrite of L5PCs but facilitate the action potential (AP) induction for proximal inputs. By using two models with different Ca^2+^ channel distributions we showed that the shunt-inhibitory effect of the *I_h_* channels requires presence of a hot zone of low-voltage activated (LVA) Ca^2+^ channels at the mid-distal apical dendrite. Furthermore, we showed that neuromodulators had a similar bimodal location-dependent effect on L5PC excitability. We also demonstrated that maximal neuromodulatory effects can be brought about by combining *I_h_*-enhancing neuromodulation at the proximal dendrite and *I_h_*-inhibiting neuromodulation at the distal dendrite, or vice versa. Our analysis uncovers the effects that neuromodulation of *I_h_* can have on L5PCs at a single-cell level. This sheds new light on how L5PCs integrate information and enable higher-order functions of the brain.

## 2 Methods

### 2.1 Neuron models

We employed two models of L5PCs: the “Hay model” [Hay et al., 2011] and the “Almog model” [Almog and Korngreen, 2014], both of which were multicompartmental Hodgkin-Huxley type of models with reconstructed layer V thick-tufted pyramidal neuron morphologies. The ionic current species of the two models are listed in Table 1. In the Hay model, the ion-channel conductances were constant along the dendrites, except for the *I_h_* channel whose conductance grows exponentially with the distance from the soma and the Ca^2+^ channels where a hot zone of Ca^2+^ channels (10× larger HVA Ca^2+^ channel conductance and 100× larger LVA Ca^2+^ channel conductance) was present at the apical dendrite at a distance from 685 to 885 μm from the soma [Hay et al., 2011]. In the Almog model, there was no hot zone of Ca^2+^ channels, but all ion-channel conductances varied spatially (usually piece-wise linear) along the dendrites [Almog and Korngreen, 2014]. In addition to describing the dynamics of these ionic channels, the models also describe the dynamics of the intracellular Ca^2+^ concentration, [Ca^2+^]_i_, which affects the currents conducted by SK and BK channels. According to the models, [Ca^2^+]_i_ is increased by the current flow through Ca^2+^ channels, and is otherwise decreased towards a resting-state level of [Ca^2^+]_i_.

**Table 1:** Current species in the Hay and Almog models.

| Hay Model | Almog model |
| --- | --- |
| Fast inactivating Na<sup>+</sup> current ($I_{Nat}$) | Fast inactivating Na<sup>+</sup> current ($I_{Nat}$) |
| Non-specific cation current ($I_h$) | Non-specific cation current ($I_h$) |
| High-voltage-activated Ca<sup>2+</sup> current ($I_{CaHVA}$) | High-voltage-activated Ca<sup>2+</sup> current ($I_{CaHVA}$) |
| Low-voltage-activated Ca<sup>2+</sup> current ($I_{CaLVA}$) | Medium-voltage-activated Ca<sup>2+</sup> current ($I_{CaMVA}$) |
| Fast inactivating K<sup>+</sup> current ($I_{Kt}$) | Fast inactivating K<sup>+</sup> current ($I_{Kt}$) |
| Slow inactivating K<sup>+</sup> current ($I_{Kp}$) | Slow inactivating K<sup>+</sup> current ($I_{Kp}$) |
| Small-conductance Ca<sup>2+</sup>-activated K<sup>+</sup> current ($I_{SK}$) | Small-conductance Ca<sup>2+</sup>-activated K<sup>+</sup> current ($I_{SK}$) |
| Fast non-inactivating K<sup>+</sup> current ($I_{Kv3.1}$) | Large-conductance voltage and Ca<sup>2+</sup>-gated K<sup>+</sup> current ($I_{BK}$) |
| Muscarinic K<sup>+</sup> current ($I_m$) | Passive leak current ($I_{leak}$) |
| Persistent Na<sup>+</sup> current ($I_{Nap}$) | |
| Passive leak current ($I_{leak}$) | |

All simulations were run using NEURON software (version 7.8.2) with adaptive time-step integration. Our simulation scripts (interfaced through Python, versions 3.7.5 and 3.9.1 tested) are publicly available at http://modeldb.yale.edu/267293 (password “modulhcn” required during peer-review).

### 2.2 Stimulation protocols

In Section 3.1, we stimulated the center of the soma with a square pulse current, starting at 200 ms and lasting until the end of the simulation (16 s). The spiking frequency was determined based on the number of spikes from 500 to 16000 ms. In Section 3.2, we stimulated dendritic sections with a short square pulse current (0.2 ms) or with a conductance-based, alpha-shaped (time constant 5 ms) glutamatergic (reversal potential *E*_glu_ = 0 mV) input. When choosing the location along the dendrite the thickest dendritic section at a given distance was selected as in [Hay et al., 2011]. In Sections 3.3–3.4, the glutamatergic synaptic inputs were modelled with more precision (except for the simulations of Fig. 5C–D), similar to [Compte et al., 2000, Aćimović et al., 2021]: The AMPAR-mediated currents were modelled as

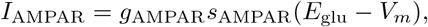

where the synaptic variable *s* increased instantaneously with incoming spikes and decayed exponentially as 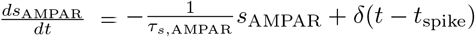 with a time constant *τ*_*s*,AMPAR_ = 2 ms. The NMDAR-mediated currents were modelled as

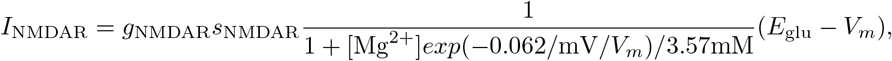

where the synaptic variables *s*_NMDAR_ and *x*_NMDAR_ obeyed the following dynamics:

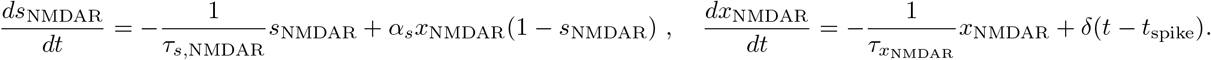

The rise time was *τ*_*x*,NMDAR_ = 2 ms and the decay time *τ*_*s*,NMDAR_ = 100 ms, and the rate of current activation was set *α_s_* = 2 kHz as in [Compte et al., 2000]. The NMDAR-mediated current was also used Fig 3E in Section 3.2, where *g*_AMPAR_ = 0 uS, whereas in Sections 3.3–3.4 the conductances *g*_AMPAR_ and *g*_NMDAR_ were set the same. In Section 3.4, we also modelled the effects of concurrent GABAergic inhibition of the basal dendrite, which was modelled the same way as AMPAR-mediated currents but the reversal potential was set −80 mV. The spike time *t*_spike_ of the AMPAR- and NMDAR-mediated inputs was the same (2000 ms) for all synapses, while the spike time of GABAergic inputs was randomly picked between 1975–2025 ms.

### 2.3 Alterations of ion-channel properties

#### Blockage of *I_h_*

We blocked the *I_h_* currents by setting the maximal conductance to 0 everywhere (unless otherwise stated) in the neuron.

#### Dopaminergic and cholinergic modulation of *I_h_*

We modelled the dopaminergic (D1-receptor-mediated) modulation of *I_h_* by increasing the half-inactivation voltage by +10 mV or +5 mV in the Hay or Almog model, respectively, and the cholinergic modulation of *I_h_* by decreasing it by the same amount [Zhao et al., 2016, Byczkowicz et al., 2019]. We used the smaller voltage difference for the Almog model due to the relatively large effect of ±10 mV shifts (data not shown).

#### Removal of the hot zone of Ca^2+^ channels from the Hay model

We used the same conductance of HVA and LVA Ca^2+^ channels in the area of the hot zone as elsewhere in the apical dendrite (*g*_CaHV_ A = 55.5 uS/cm^2^, *g*_CaLV_ A = 187 uS/cm^2^).

#### Addition of a hot zone of Ca ^2+^ channels to the Almog model

We adopted the LVA Ca^2+^ current from the Hay model in addition to the HVA and MVA Ca^2+^ currents native to the Almog model. We added LVA Ca^2+^ channels to the apical dendrite with a maximal conductance of 300 mS/cm^2^ for dendritic sections with a distance of 585–985 μm to the soma and 3 mS/cm^2^ for the others.

## 3 Results

### 3.1 Activation of *I_h_* increases L5PC activity when stimulated at soma

*I_h_* blockage has been shown, both experimentally (e.g., by application of ZD7288 [Maccaferri and McBain, 1996, Endo et al., 2008]) and computationally [Kase and Imoto, 2012], to lead to decreased neuron spiking. Here, we replicated this result with our biophysically detailed neuron models of a L5PC neuron, namely, the Almog model (Fig. 1A) [Almog and Korngreen, 2014] and the Hay model (Fig. 1B) [Hay et al., 2011], by considering the f-I curve of the L5PC under various operations on the *I_h_* channel. A block of *I_h_* decreased the rate of firing in response to DC applied to soma, both in the Almog and Hay models, although the effects were considerably larger in the Almog model (Fig. 1C, blue) than in the Hay model (Fig. 1D, blue). Accordingly, an increase in *I_h_* conductances increased the firing rates in both models (Fig. 1C–D, red curves). Likewise, a model of neuron-wide cholinergic modulation (half-inactivation voltage decreased) decreased the firing rates, and a dopaminergic modulation (half-inactivation voltage augmented) increased the firing rates in both models (Fig. 1E–F). This was also the case when *I_h_* currents were blocked or enhanced only at the dendrites (Fig. S1A–D) instead of both soma and dendrites: both *I_h_* blockage (Fig. S1A–B, blue) and cholinergic modulation (Fig. S1C–D, blue) at the dendrites decreased the L5PC firing rates, while *I_h_* over-expression (Fig. S1A–B, red) and dopaminergic modulation (Fig. S1C–D, red) at the dendrites increased the firing rates. Taken together, these results support the role of *I_h_* current as an enhancer of L5PC activity when the neuron is stimulated at the soma.

**Figure 1:**
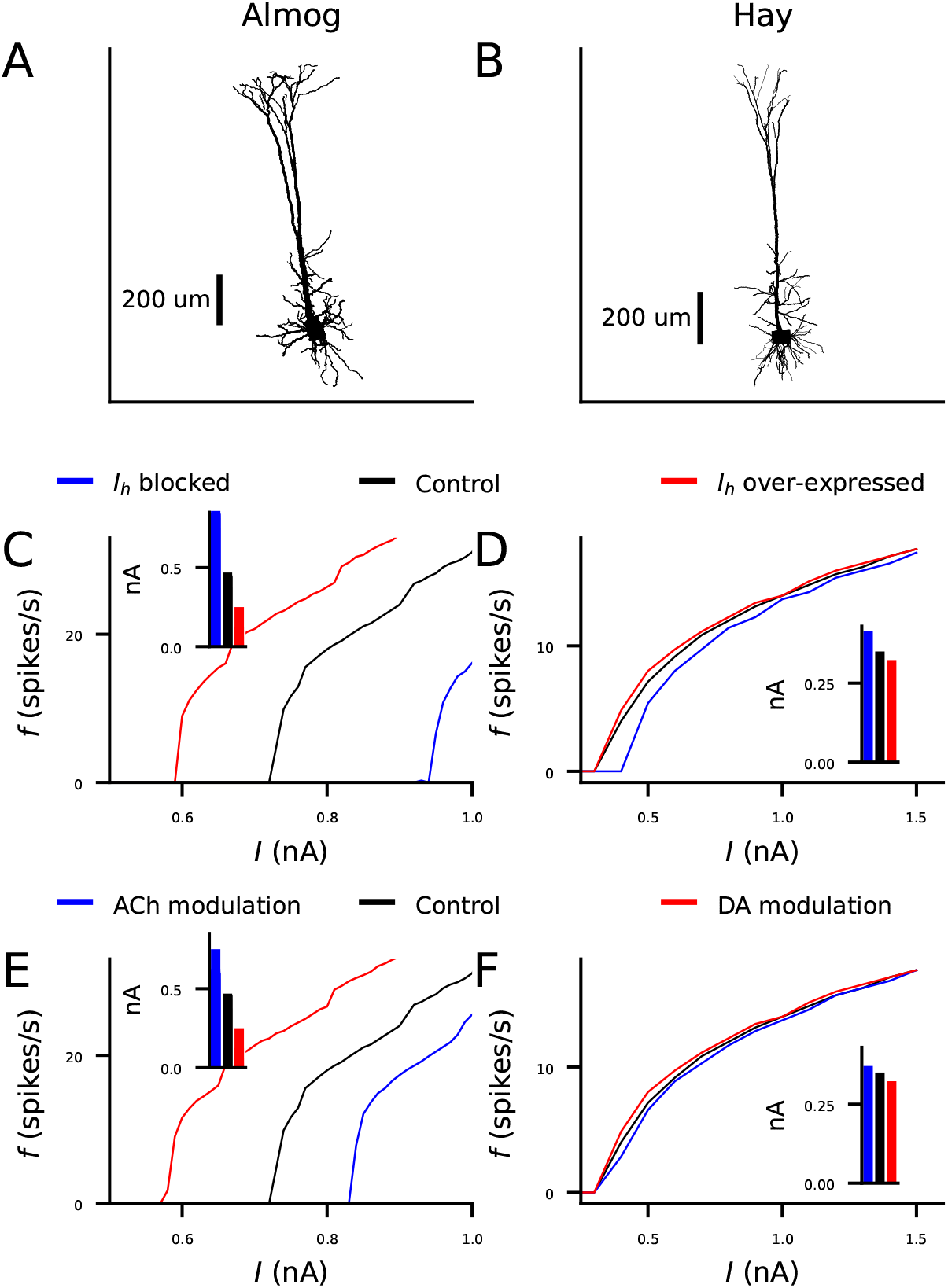
*I_h_* activation increases the frequency of action potentials in response to somatic DC in L5PCs. **A–B**: The morphology of the Almog (A) and Hay (B) moadel neurons. **C–D**: The frequency of APs (y-axis) in response to somatic DC of a given amplitude (x-axis) in Almog (C) and Hay (D) model neurons under up- or down-regulated *I_h_* channels. Black: control neuron. Blue: *I_h_* conductance blocked. Red: *I_h_* conductance increased by 100%. **E–F**: The frequency of APs response to somatic DC in Almog (E) and Hay (F) model neurons under different neuromodulatory states. Black: control neuron. Blue: cholinergic neuromodulation, modelled as a −5 mV (E) or −10 mV (F) shift in half-inactivation potential of the *I_h_* channels. Red: dopaminergic neuromodulation, modelled as a +5 mV (E) or +10 mV (F) shift in half-inactivation potential of the *I_h_* channels. The insets of panels (C)–(F) show the threshold current amplitudes for inducing an AP (note that these can be different from the onsets of the f-I curves since some DC amplitudes only cause one spike).

### 3.2 *I_h_* activation can increase AP threshold in response to apical dendritic stimulation in an L5PC when a hot zone of Ca^2+^ channels is present

The reversal potential of the *I_h_* typically lies in the range −45 – −30 mV, which grants the *I_h_* channel the possibility to shunt, that is, to inhibit inputs that would otherwise depolarize the membrane to potentials higher than this. Indeed, in [Poolos et al., 2002], increased *I_h_* activation by lamotrigine application led to decreased firing response to dendritic stimuli in a CA1 neuron, and they reproduced their findings with a computational model consisting of a realistic morphology and one type of sodium and two types of potassium channels. In line with this, in [Magee, 1999], pharmacological blockage of *I_h_* increased the amplitude of distally elicited EPSPs compared to proximally elicited ones in a CA1 neuron. Computational models have suggested that the observed phenomena could be caused by secondary mechanisms, where a blockage or altered expression of *I_h_* channels also indirectly affects conductance of other ion channels, such as Twik-related acid-sensitive K^+^ (TASK) channels [Migliore and Migliore, 2012, Kelley et al., 2021]. Here, focusing solely on L5PCs, we explored by means of computational modelling the possibility that the inhibitory actions attributed to *I_h_* activity are caused by direct shunting effects of the current without concurrent changes in the conductance of other ion channels.

We simulated the injection of a strong dendritic square-pulse current stimulus of 0.2 ms duration that locally depolarizes the dendrite. We measured the somatic response in the presence of *I_h_* current and compared it to the response in absence or partial absence of the current. We varied the site of dendritic stimulation from 50 to 1000 μm with an interval of 50 μm and used the bisection method to find the AP threshold for each stimulation site.

The Almog model neuron consistently predicted that the presence of *I_h_* current facilitated the AP initiation by a dendritic stimulus (Fig. 2A–D). Blocking the *I_h_* currents hyperpolarized the basal membrane potential and increased the threshold current for inducing a spike with a dendritic stimulus at a distance of 500 μm from the soma (Fig. 2A). The threshold current was increased in the Almog model across the apical dendrite (Fig. 2B). For stimulation sites further than 400–500 μm from the soma the threshold currents implied unrealistically high (> 100 mV) membrane potentials at the site of stimulation (Fig. 2C). We thus confirmed our results using conductance-based inputs: whenever a spike could be initiated by an alpha-shaped glutamatergic synaptic conductance, the threshold conductance was increased in the *I_h_*-blocked model compared to the control Almog model (Fig. 2D).

**Figure 2:**
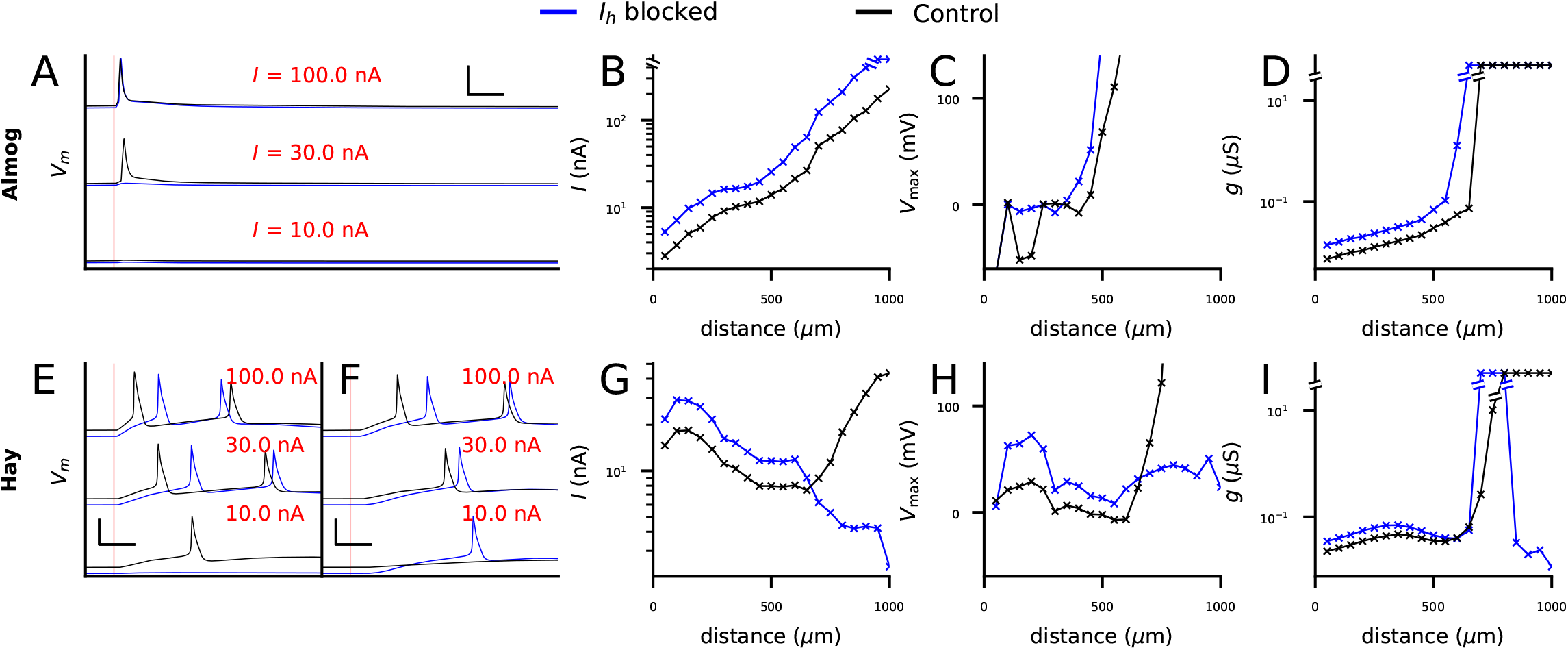
*I_h_* activation may increase or decrease the threshold for L5PC action potential firing by a strong apical dendritic input, depending on the location of the input. **A**: Somatic membrane potential time courses according to the Almog model in response to 2-ms square-pulse current with an amplitude of 100 (top), 30 (middle) or 10 nA (bottom), injected at the apical dendrite 500 μm from the soma. For control neuron (black), both 30 and 100 nA stimuli induced a spike, while for *I_h_*-blocked neuron (blue), 100 nA stimulus induced a spike while 30 nA stimulus did not. **B**: Threshold current amplitudes for 2-ms square-pulse inputs at the apical dendrite at different distances from the soma. Black: Almog-model control neuron, blue: Almog-model neuron with *I_h_* blockage. **C**: Peak membrane potential at the site of current injections, given the threshold-current of (B). **D**: Threshold conductances for an alpha-shaped glutamatergic input with time constant 3 ms. Black: Almog-model control neuron, blue: Almog-model neuron with *I_h_* blockage. **E–F**: Somatic membrane potential time courses according to the Hay model in response to 2-ms square-pulse current with an amplitude of 100 (top), 30 (middle) or 10 nA (bottom), injected at the apical dendrite 500 (E) or 800 (F) μm from the soma. For a stimulus 500 μm from the soma (E), 10.0 nA stimulus induced a spike in the control neuron (black) but not in the *I_h_*-blocked neuron (blue), while at 800 μm this was reversed. **G–I**: The experiments of (B–D) repeated for Hay model.

In contrast to the Almog model predictions, the Hay model predicted that blockage of *I_h_* current may either facilitate or hinder the AP initiation by dendritic stimulation, depending on the distance from the soma (Fig. 2E–I). Although the blockage of *I_h_* currents resulted in hyperpolarization of the baseline membrane potential at a distance of both 500 and 800 μm from the soma (Fig. 2E–F), the threshold current was increased for stimulus at 500 μm (Fig. 2E) and decreased at 800 μm (Fig. 2F) from the soma. The threshold currents applied to dendritic sites further than 700 μm from the soma in the presence of *I_h_* currents caused unrealistically high local membrane potentials (Fig. 2H), and thus we replicated the result of Fig. 2G using conductance-based stimuli (Fig. 2I): the threshold conductance was larger for the *I_h_*-blocked than the default Hay model when the stimulus site was closer than 800 μm from the soma, and vice versa for stimuli further than 800 μm (Fig. 2I). The switch of *I_h_* becoming inhibitory happened around 650–850 μm (Fig. 2G,I), which coincides with the hot zone of Ca^2+^ channels in the Hay model (685–885 um) [Hay et al., 2011].

We next analyzed the contributions of the Ca^2+^ channels to the switch in threshold currents between control and *I_h_*-blocked neuron models. Complete blockage of LVA Ca^2+^ channels radically increased the threshold currents in the *I_h_*-blocked Hay-model neuron, abolishing the switch in threshold currents (Fig. 3A), while blocking any of the other voltage-gated ion channels (see Table 1) from the Hay model did not change the qualitative behaviour of Fig. 2G (Fig. S2). The relatively small effects of blockade of HVA Ca^2+^ channels and SK channels was surprising in light of our previous computational studies highlighting the role of these channels in shaping L5PC activity [Mäki-Marttunen et al., 2017, Mäki-Marttunen et al., 2019]. In fact, it was sufficient to only remove the excessive LVA Ca^2+^ channels from the hot zone of Ca^2+^ channels: when the same LVA Ca^2+^ channel conductances were used in the hot zone as in the rest of the apical dendrite while other ion-channel conductances were untouched, *I_h_* current blockage always led to an increased threshold current, similar to the Almog model (Fig. 3B) — we also replicated this result with conductance-based inputs (Fig. 3C). As suggested by previous modelling work of CA1 neurons [Tsay et al., 2007], the reason behind the interaction of LVA Ca^2+^ currents and the *I_h_* current was that in the presence of *I_h_* currents, the LVA Ca^2+^ channels were highly inactivated at resting membrane potential (Fig. 3D). We confirmed the decisive role of the LVA Ca^2+^ channel inactivation with simulations of an isolated Hay-model compartment from the distal apical dendrite: when the time course of *activation* variable of the LVA Ca^2+^ channels was artificially replaced by the corresponding time course recorded in the absence of *I_h_* channels, the *I_h_*-blocked dendrite remained more excitable than the dendrite with *I_h_* channels intact, but when the time course of *inactivation* variable was replaced by that recorded in the absence of *I_h_* channels, the shunting effect disappeared (Fig. S3). In line with these observations, when a hot zone of LVA Ca^2+^ channels was added to the Almog model, we observed a qualitatively similar switch of threshold current amplitudes between *I_h_*-blocked and control case (Fig. 3E). Taken together, our simulations suggest *I_h_* channel activity can increase the threshold current in distal dendritic stimuli (i.e., it can have a shunting inhibitory effect on excitatory inputs) when the apical trunk expresses LVA Ca^2+^ channels.

**Figure 3:**
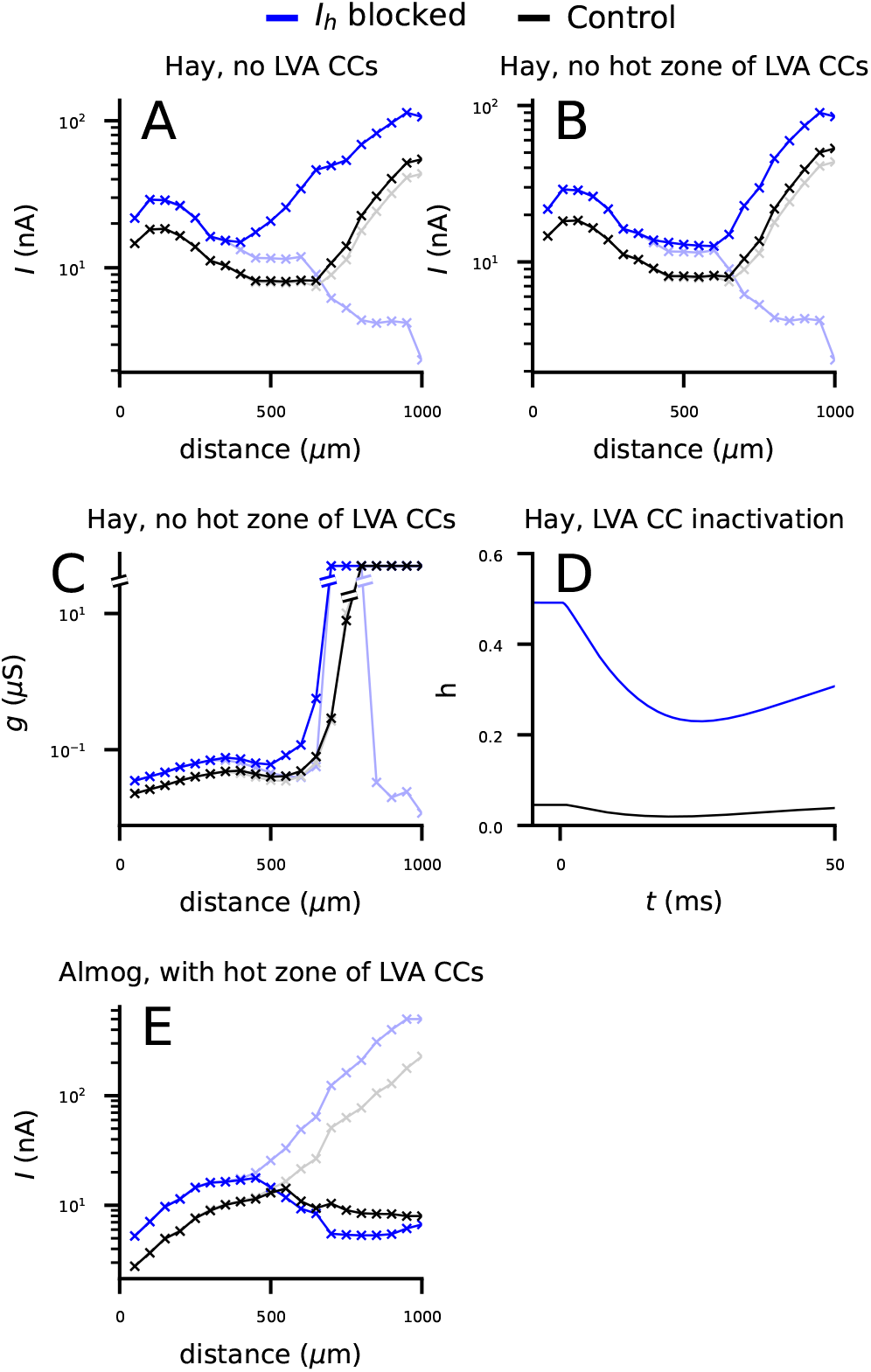
The shunting inhibitory effect of *I_h_* channels is mediated by Ca^2+^ channels in the apical dendrite. **A**: Threshold current amplitudes for 2-ms square-pulse inputs at the apical dendrite according to Hay model without LVA Ca^2+^ channels. Black: Hay-model neuron with LVA Ca^2+^ channel blockage, blue: Hay-model neuron with LVA Ca^2+^ and *I_h_* channel blockage. Dim curves: the data from Fig. 2G where LVA Ca^2+^ channels were intact. **B**: Threshold current amplitudes according to Hay model without a hot zone of LVA Ca^2+^ channels with (black) or without (blue) *I_h_* channels. Dim curves: the data from Fig. 2G where LVA Ca^2+^ channels had higher conductance in the hot zone (650–850 μm from the soma). **C**: Threshold conductances for an alpha-shaped inputs from Fig. 2D in a Hay-model neuron without a hot zone of LVA Ca^2+^ channels with (black) or without (blue) *I_h_* channels. Dim curves: the corresponding data from a Hay model where LVA Ca^2+^ channels were intact. **D**: The time course of the inactivation variable of *I_h_* according to the Hay model in response to an alpha-shaped conductance (onset at t=0 ms) of amplitude 0.01 nS. **E**: Threshold current amplitudes in an Almog-model neuron where MVA Ca^2+^ channels in the apical dendrite were replaced by LVA Ca^2+^ channels with conductance 0.003 S/cm^2^, except for apical dendritic sections at a distance of 585–985 μm from soma where the conductance was 0.3 S/cm^2^, with (black) or without (blue) *I_h_* channels. Dim curves: the data from Fig. 2B where the Ca^2+^ channels were as in the default Almog model.

### 3.3 Shunting inhibition by *I_h_* current can also occur for spatially distributed stimuli

Until now, we have stimulated the neuron with a single input in the soma or in the dendrite at a time, while it is expected that many excitatory synaptic inputs are needed to induce an AP. For this reason, we were not always able to initiate APs by stimulating distal dendrites without using unrealistically strong inputs (Fig. 2C–D,H–I). We thus went on to explore the effects of *I_h_* channel activity on AP induction in L5PCs when the stimuli arrive simultaneously at different locations of the dendrite. To do this, we injected AMPAR- and NMDAR-mediated glutamatergic synaptic currents into randomly picked locations at a given distance from the soma. For each set of synapse locations, we searched for the threshold conductance for inducing a somatic AP. We repeated the procedure *N_samp_*=40 times to obtain distributions of threshold conductances, which we used for statistical tests between *I_h_*-blocked and control neurons.

We first distributed the synapses all along the apical (Fig. 4A, cyan to blue) or basal (Fig. 4A, red to orange) dendrite and activated them simultaneously. In both of these cases, presence of *I_h_* currents lowered the AP threshold in both the Almog (Fig. 4B–C) and Hay (Fig. 4D–E) models. This was expected, since we observed similar trend in the f-I curves for somatic stimulus (Fig. 1C-D) and in the threshold currents for proximal inputs at the apical dendrite (Fig. 2B,G) — and although the distal inputs showed an opposite trend in the Hay model, the proximal inputs are likely to be more determinant for AP initiation than distal inputs of the same strength. We next distributed the synapses on the apical dendrite at intervals [*x*_1_, *x*_2_] from soma where we varied *x*_1_ and *x*_2_ from 200 to 1300 μm (furthest point of the apical dendrite) in intervals of 100 μm. In the Almog model, *I_h_* activity always facilitated the neuron firing (Fig. 4F), while in the Hay model, *I_h_* activity facilitated the neuron firing for proximal inputs and raised the threshold for distal inputs (Fig. 4G). Apart from the most distal parts of the Almog-model neuron (Fig. 4F), APs could be initiated with physiologically realistic (conductance of a single synapse < 1 nS) stimulation of all parts of the dendritic trees of the model neurons. Despite the variability in threshold conductance depending on the exact location of the synaptic inputs at the given locations, all differences between *I_h_*-blocked and control neurons were statistically significant (U-test, p<0.05; Bonferroni corrected by the number (66) of statistical tests), except for the ones where no AP was initiated for any tested synaptic conductance either in the control or *I_h_*-blocked neuron (grey squares in the lower right triangle of Fig. 4F). This applied also to the Almog model supplemented with densely distributed LVA Ca^2+^ channels to form a hot zone of Ca^2+^ channels in the apical dendrite (Fig. S4A). Taken together, our results suggest that spatially distributed stimuli with physiologically realistic conductances may be shunted by *I_h_* channels in presence of a hot zone of LVA Ca^2+^ channels, but without the hot zone the *I_h_* channels only contribute to lowering the AP threshold of spatially distributed stimuli.

**Figure 4:**
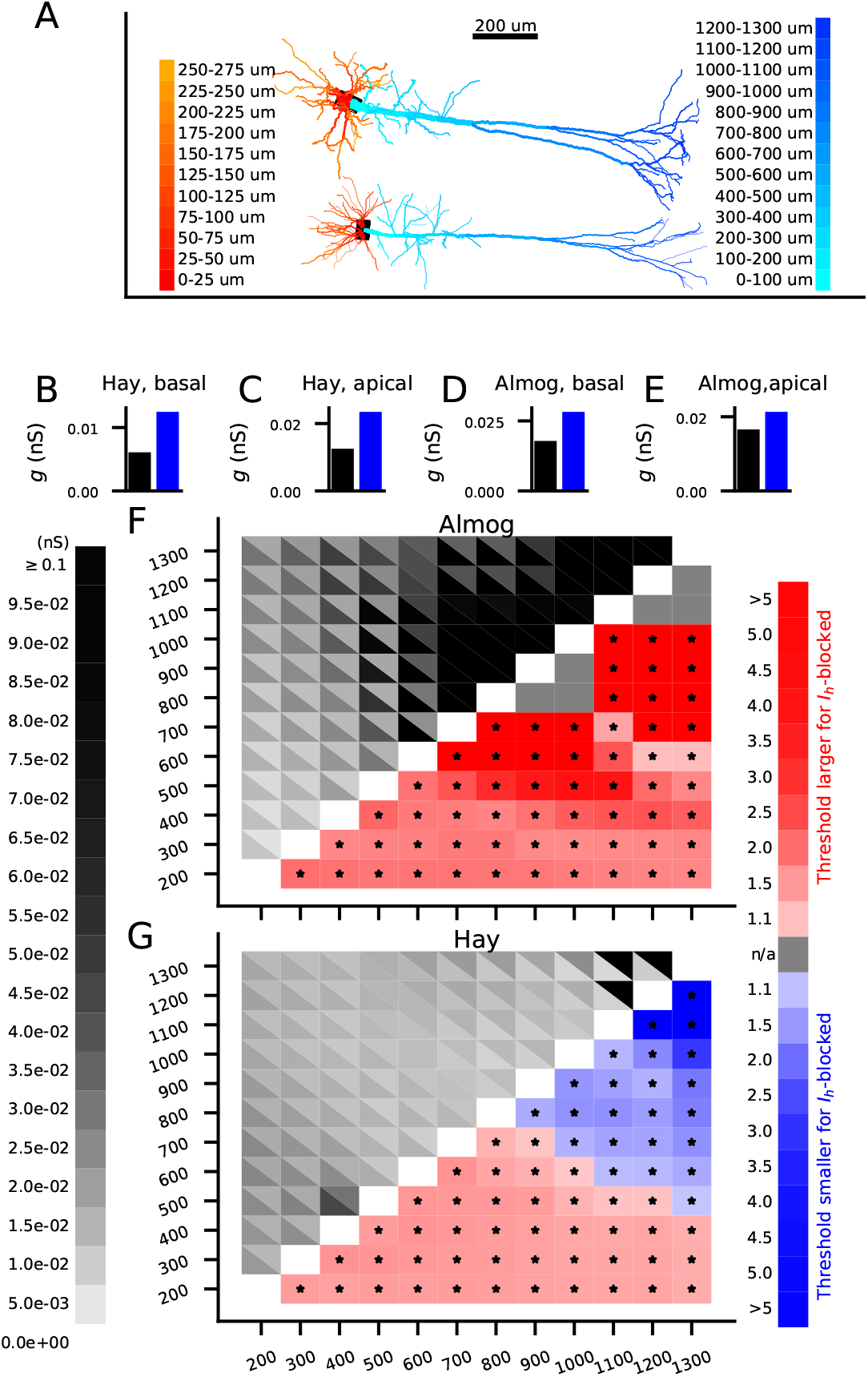
*I_h_* channels can shunt spatially distributed excitatory synaptic inputs when the inputs arrive at the distal dendrite. **A**: Almog- (top) and Hay-model (bottom) morphology color-coded according to distance from soma. **B–E**: Threshold conductance for a set of 2000 excitatory synapses (simultaneously activated) to induce an AP in the Almog (B–C) or Hay (D–E) model in a control (black) or *I_h_*-blocked (blue) L5PC. In (B) and (D), the synapses were uniformly distributed along the basal dendrite, whereas in (C) and (E), the synapses were uniformly distributed along the apical dendrite. **F**: *Upper left grid*: The threshold conductances for a set of 2000 excitatory synapses (simultaneously activated) to induce an AP in the Almog model. The synapses were uniformly distributed along the apical dendrite at distances [*x*_1_, *x*_2_] from the soma where the parameters *x*_1_ (x-axis) and *x*_2_ (y-axis) ranged from 200 μm to 1300 μm in intervals or 100 μm. In each grid slot, the color of the upper right triangle indicates the threshold conductance in the control neuron wherease that of the lower left triangle indicates the threshold conductance in the *I_h_*-blocked neuron. *Lower right grid*: The factor by which the threshold conductance of the *I_h_*-blocked neuron is larger (red) or smaller (blue) than that of the control neuron for the stimuli distributed in sections ranging from 200–1200 μm (y-axis) to 300–1300 μm (x-axis). Asterisks indicate statistically significant differences (U-test, p<0.05/66). Grey squares represent sections where the set of stimuli was unable to induce an AP for all tested conductances (until 0.1 pS) in both *I_h_*-blocked and control L5PC. **G**: The experiment of (F) repeated for the Hay model.

### 3.4 Dopaminergic modulation of distal dendrites and cholinergic modulation of proximal dendrites as well as GABAergic inhibition of the basal dendrites strengthen the shunt-inhibitory role of *I_h_* channels

The above analyses highlighted the bimodal effect of *I_h_* currents when the neuron was uniformly affected by *I_h_* blocker. However, pyramidal neurons express a large set of neurotransmitter receptors that are non-uniformly distributed or selectively activated by presynaptic connections. Here, we explored how the shunt/excitation dichotomy of the *I_h_* channels is affected by interaction of different neurontransmitter systems in different parts of the dendritic tree.

First, we replicated the qualitative result of Fig. 4G using whole-cell neuromodulation — i.e., we used cholinergic modulation of *I_h_* instead of *I_h_* blockage and dopaminergic modulation as an *I_h_*-enhancer. As expected, the Hay model predicted that whole-cell cholinergic modulation of *I_h_* channels increased the threshold for distal inputs and decreased the threshold for proximal inputs, while dopaminergic modulation had the opposite effect (Fig. S4B–C).

We next quantified the effect of *I_h_* activity on the threshold conductance of apical stimuli in presence of synaptic inputs arriving at the basal dendrite. When glutamatergic stimulation was applied at the same time to both basal and apical dendrite of the Hay-model neuron, *I_h_* current had a mostly excitatory effect except for the stimuli applied to the very distal parts of the apical dendrite (Fig. 5A). By contrast, when GABAergic stimulation was applied to the basal dendrite within a window of ±25 ms from the glutamatergic stimulation at the apical dendrite, we observed a stronger shunting inhibition effect of the *I_h_* current (Fig. 5B). We repeated these simulations using simpler, current-based analyses (Fig. 5C–D). Our model of cholinergic neuromodulation had similar effects: neuron-wide cholinergic modulation in presence of glutamatergic inputs at the basal dendrite increased the AP threshold only for very distal apical stimuli, while in the presence of GABAergic stimulation it increased the AP threshold also for middle-apical (600–700 μm) inputs (Fig. S5). These results suggest that simultaneous activation of excitatory synapses at basal and apical dendrites constrain the shunt-inhibition effect of *I_h_* channels to the most distal parts of the apical dendrite.

**Figure 5:**
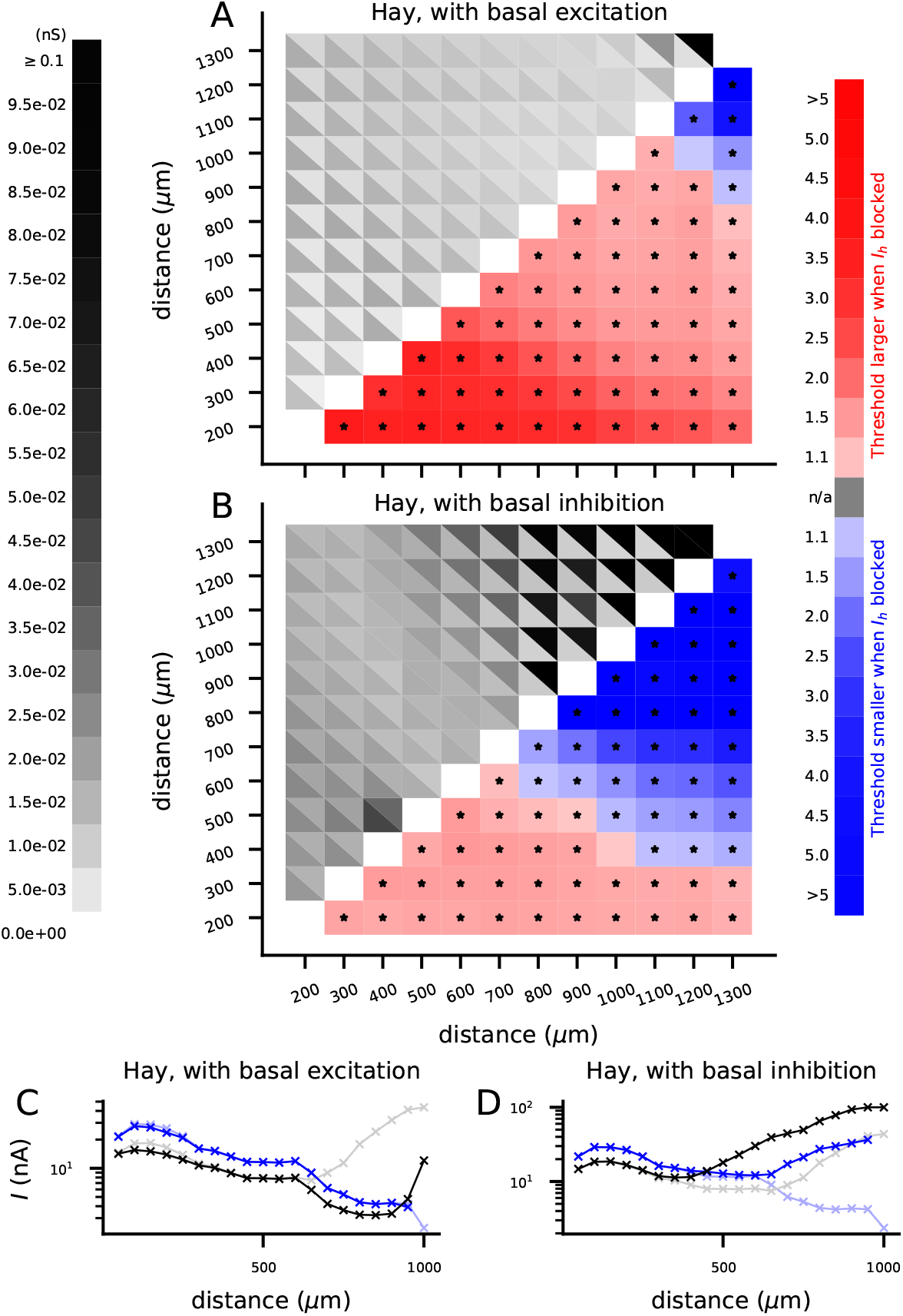
The shunting effect of *I_h_* is constrained to distal parts of apical dendrite by simultaneous glutamatergic stimulation and strengthened by GABAergic stimulation of the basal dendrite. **A–B**: The Hay-model predictions for the AP threshold for spatially distributed apical dendritic stimulation in presence and absence of *I_h_* channels when the basal dendrite is simultaneously stimulated with glutamatergic (A) or GABAergic (B) inputs. See Fig. 4G for details. For the glutamatergic stimulation of the basal dendrite, we distributed 2000 glutamatergic synapses across the basal dendritic tree, and their conductance was set 80% of the AP threshold conductance in the control condition (18 pS, see Fig. 4D). For the GABAergic stimulation of the basal dendrite, we distributed 2000 GABAergic synapses across the basal dendritic tree, and we randomly picked their activation times from the uniform distribution ±25 ms from the apical activation time and set their conductance 200 pS. **C–D**: The AP threshold for a single current-based apical dendritic stimulus in presence (black) and absence (blue) of *I_h_* channels when the basal dendrite is simultaneously stimulated with glutamatergic (C) or GABAergic (D) inputs. The dim curves show the data from Fig. 2G where there was no stimulation of the basal dendrite. In (C), we stimulated the basal dendritic compartment at 50 μm from the soma with a conductance-based, alpha-shaped glutamatergic (reversal potential 0 mV) input whose amplitude was 100 nS, and in (D) with a GABAergic (reversal potential −80 mV) input with amplitude 5000 nS.

Finally, we analyzed the effects of a partial neuromodulation of the apical dendrite and interactions of cholinergic and dopaminergic neuromodulation in affecting the AP threshold in the apical dendrite. When proximal (up to 500 μm from the soma) apical dendrite of the Hay model was modulated by dopamine (Fig. 6A, blue) or acetylcholine (Fig. 6B, blue), the AP threshold for a set of simultaneously activated glutamatergic synapses was lowered or increased, respectively, across the stimulus locations. In this experiment, we distributed the synapses across dendritic locations at a distance [*x*_1_,*x*_1_+100μm] from the soma, where we varied *x*_1_ from 200 to 1200 μm. When the modulation of the proximal apical dendrite by dopamine was accompanied by cholinergic modulation of the distal (from 500 μm from the soma onwards) apical dendrite, the neuron was made yet more excitable at the distal dendrites (Fig. 6A, magenta) — and accordingly, when the cholinergic modulation of the proximal apical dendrite was accompanied by dopaminergic modulation of the distal apical dendrite, the cell was yet less excitable at the distal apical dendrite (Fig. 6B, magenta). When we repeated the experiment using wider distributions of synapses as done in the experiments of Fig. 4F–G and 5A–B, a similar result was obtained (Fig. 6C–D). We also performed a similar experiment using single current-based stimuli along the apical dendrite and different L5PC models. In these experiments, dopaminergic (cholinergic) modulation of proximal apical *I_h_* channels lowered (raised) the AP threshold of a single current-based stimulus, both in the Hay and Almog models and in the Almog model expressing a hot zone of LVA Ca^2+^ channels (Fig. S6, blue). Moreover, this modulation in combination of cholinergic (dopaminergic) modulation of the distal dendrite further lowered (raised) the AP threshold in the distal apical dendrite (Fig. S6, magenta) — this applied to the Hay model (Fig. S6A,D) and the Almog model with the hot zone of Ca^2+^ channels (Fig. S6C,F), but the native Almog model predicted in-between levels of excitability for neuromodulator combinations (Fig. S6B,E). Likewise, the combination of the opposite modulations at proximal and distal dendrite was more effective than the modulation of distal apical dendrite alone (data not shown). Taken together, our modelling results suggest that *I_h_* channel-activating neuromodulation in tha proximal apical dendrite increases the L5PC excitability, and that in L5PCs expressing a hot zone of LVA Ca^2+^ channels, the neuron is made yet more excitable by *I_h_*-suppressing neuromodulation in the distal apical dendrite.

**Figure 6:**
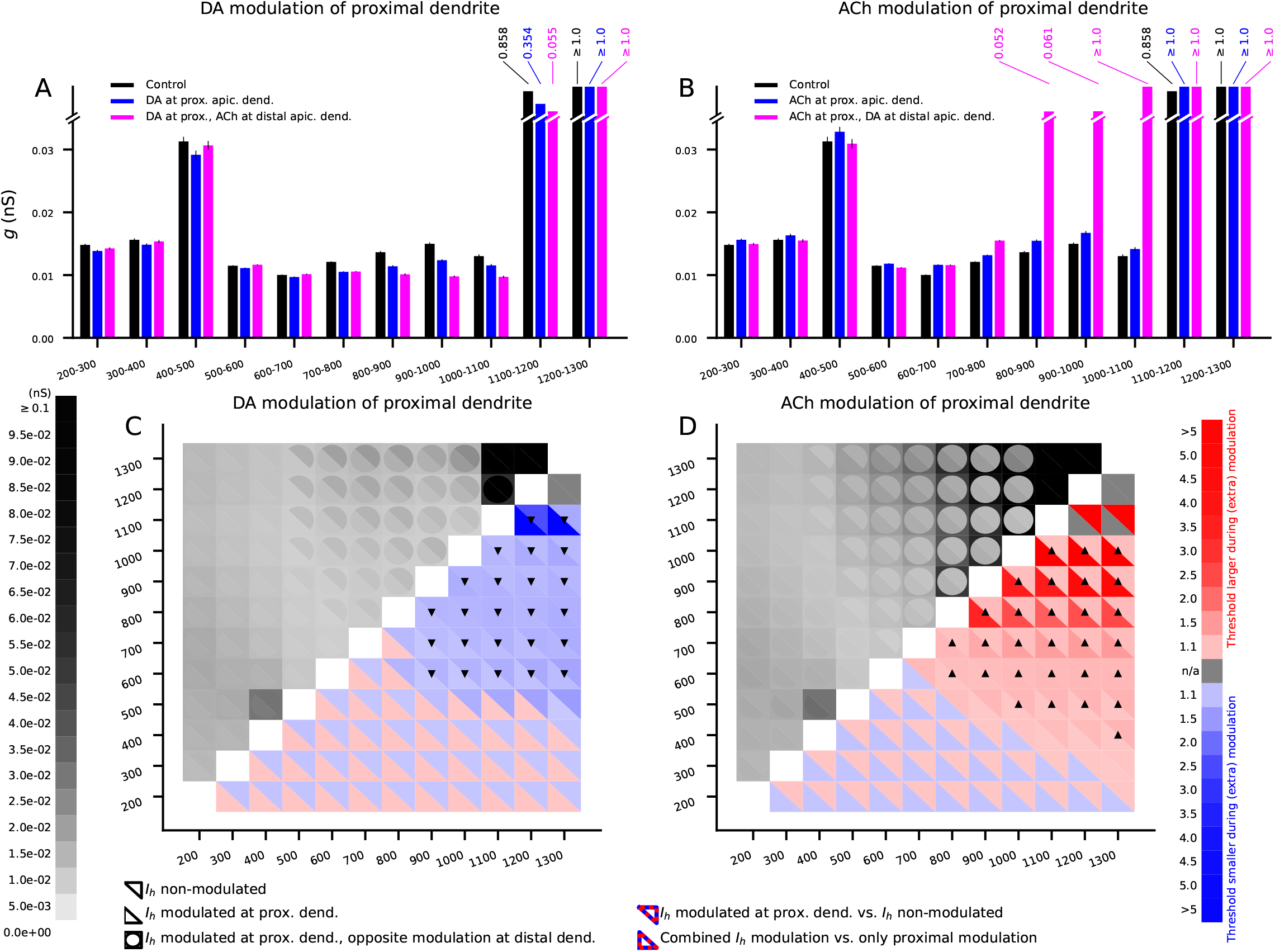
Combination of dopaminergic and cholinergic neuromodulation can increase or decrease the AP threshold throughout the apical dendrite in the Hay-model L5PC. **A**: The threshold conductances for a set of 2000 excitatory synapses (distributed across apical dendrite at distances [*x*_1_,*x*_1_+100 μm] from the soma, simultaneously activated) to induce an AP in the Hay model. Black: Control Hay model. Blue: Dopaminergic modulation of the *I_h_* channels in proximal apical dendrite (where the center of the compartment was located nearer than 500 μm to the soma). Magenta: Dopaminergic modulation of proximal apical dendrite and cholinergic modulation of the distal apical dendrite. **B**: The experiment of (A) repeated with the opposite modulation, i.e., cholinergic modulation of proximal apical dendrite and dopaminergic modulation of the distal apical dendrite. **C**: The experiment of (A) repeated by varying the width of the synapse distribution as in Fig. 4F–G. *Upper left grid*: In each slot, the color of the upper right triangle indicates the threshold conductance in the control neuron, the lower left triangle indicates the threshold when the proximal apical dendrite was modulated by dopamine, and the color of the circular frame surrounding the slot indicates the threshold when the proximal apical dendrite was modulated by dopamine and distal apical dendrite by acetylcholine. *Lower right grid*: The factor by which the threshold conductance of the modulated neuron was larger (red) or smaller (blue) than that of the less modulated neuron. In each slot, the color of the upper right triangle indicates the comparison between control neuron and proximally modulated neuron, and the lower left triangle indicates the comparison between proximally modulated neurons with and without cholinergic modulation of the distal apical dendrite. The markers denote series of statistically significant differences (U-test, p<0.05/198): Upward (Δ) or downward (▽) triangle markers mean that the threshold for proximally modulated neuron was significantly larger or smaller than that of the control neuron and the threshold for proximally (dopamine) and distally (acetylcholine) modulated neuron was larger or smaller than that of the neuron that was only proximally modulated, respectively. **D**: The experiment of (C) repeated with the opposite modulation.

## 4 Discussion

An intriguing property of *I_h_* channels is that they can exert opposite effects on the excitability of the L5PC. Previous research has explored several explanations, and point to a complex interaction of several factors, such as site of stimulation or input strength. However, experimental work is in general limited in the range of variables that can be manipulated simultaneously. To overcome these limitations, we used biophysically detailed computational modelling. The results from two different single-neuron models converged in an explanation to the differential modulation of *I_h_* on L5PC excitability: the presence of a hot zone of Ca^2+^ channels determines how the site of incoming stimulation will interact with *I_h_* currents. Our study helps explain the great flexibility that these channels confer to the pyramidal neurons. Such a flexibility is crucial for higher order brain functions needed for cognitive processing [Shine et al., 2019]. Furthermore, a disbalance in the modulatory role that *I_h_* channels has on L5PC may explain abnormal cortical processing in disorders whose symptomatology is hard to account for by simple or linear reductions in excitatory capacity.

Apical and distal dendrites of L5PCs provide distinct input to the neuron: while distal areas receive input from feedback, thalamic, and subcortical sources, proximal areas receive feed-forward inputs. In turn, these sites are themselves compartmentalized. *I_h_* channels are distributed unequally along the apical dendrite, with increasing density towards the apical tuft in the distal extreme, and therefore have a specific role in modulating the inputs that arrive at different points of the dendrite. We first studied how *I_h_* channels can modulate whether and how inputs from these different regions will be integrated in the soma. In the present work, we applied on one hand the widely used Hay model, which was developed to reproduce a wide range of L5PC features measured in a population of neurons [Hay et al., 2011], and on the other the Almog model, which was built to closely reproduce the features of a single L5PC using a parameter peeling approach [Almog and Korngreen, 2014]. Both models gave largely consistent findings, which speaks for the robustness of the found effects. By simulating excitatory inputs at increasing distance from the soma, we found that distance of the site of stimulation to the soma determines the effect of *I_h_* current on cell firing: *I_h_* current is excitatory if the stimulus is applied to the proximal apical or basal dendrite and inhibitory if the stimulus is applied to the distal apical dendrite. This is consistent with experimental work, where HCN channels were found to have inhibitory effects at the distal apical dendrite, but excitatory effects at the proximal sites (Harnett et al. 2015). A distance-dependent shunting effect of *I_h_* was also found in CA1 cells [Tsay et al., 2007]. Furthermore, we observed that the reversal point between inhibitory and excitatory effects occurred at a specific distance from the soma, which could not be explained solely by the increase in density of *I_h_* channels with distance to the soma.

We seeked to further disentangle the reason behind the effect of inputs’ distance to the soma on *I_h_* modulation. We observed that the distance effect in presence of *I_h_* current suffered a reversal around the site corresponding to the hot zone of Ca^2+^ channels. The Hay model including T-type Ca^2+^ channels, and a tuned Almog model where we added a hot zone of Ca^2+^ channels, allowed us to study the effect of removing this hot zone. Our simulations with the Hay model and the tuned Almog model agreed on *I_h_* having a purely excitatory effect in the absence of a hot zone of LVA Ca^2+^ channels in the apical dendrite, while in the presence of the hot zone, *I_h_* currents increased the AP threshold for distal apical dendritic stimuli. Our results are thus in line with [Tsay et al., 2007], where blockage of *I_h_* (both ZD7288 and -/- knockout of HCN1 were experimentally tested and computationally modelled) from CA1 neurons resulted in larger distal dendritic Ca^2+^ events, but unlike the study of [Tsay et al., 2007], our results additionally suggest a clear AP facilitating role for *I_h_* channels in the proximal dendrite regardless of the presence or absence of T-type Ca^2+^ channels. Altogether, our results suggest that *I_h_* activity mostly contributes to higher L5PC excitability but that *I_h_* channels can also shunt depolarizing inputs at the distal apical dendrite when the apical dendrite expresses strong LVA Ca^2+^ channels.

### 4.1 Neuromodulation of *I_h_* channels in L5PCs and its effects on excitability

The understanding of L5PC neuromodulation has increased during the past years although much remains to be revealed [Radnikow and Feldmeyer, 2018]. Cholinergic M1 receptors are expressed in the soma and proximal dendrites of L5PCs [Mrzljak et al., 1993, Shiozaki et al., 2001], and they are likely to be present also in more distal parts of the apical dendrite [Oda et al., 2018]. Cortical pyramidal cells of all layers also express D1-type receptors (i.e., dopamine receptors that increase cAMP levels upon activation) in their somata and apical dendrites [Bergson et al., 1995]. Likewise, cortical pyramidal cells express *β*_2_-adrenergic receptors in layers II/III as well as V [Zhou et al., 2013]. While there is a lack of data on the effects of neuromodulators on the gating properties of many ion channels that regulate L5PC excitability, the voltage-dependence of *I_h_* channels is known to be shifted toward more hyperpolarized potentials (channels more likely closed) by acetylcholine [Zhao et al., 2016] and toward depolarized potentials (channels more likely open) by cAMP-enhancing (i.e., Gs-activating) neuromodulators [Byczkowicz et al., 2019], such as dopamine-1 and *β* family of adrenoreceptors [Xing et al., 2016]. Based on these data, activation of receptors of these neuromodulatory systems were simulated by a shift of the voltage dependence of *I_h_* inactivation by 5–10 mV toward negative (acetylcholine) or positive (dopamine or norepinephrine) potentials. We note that the catecholamines dopamine and norepinephrine may exert opposite effects as well. D2-type receptors were found in a subclass of prefrontal cortical L5PCs projecting to subcortical areas [Gee et al., 2012] — these receptors are coupled to Gi/o proteins, which, opposite to Gs proteins, inhibit the cAMP production. Likewise, activation of *β*_2_-adrenergic receptors that canonically couple with Gs proteins also activate Gi/o proteins [Vasquez and Lewis, 2003]. As for cholinergic receptors, cortical pyramidal cells also express M2 receptors, which are coupled to Gi/o proteins [Mrzljak et al., 1998]. It remains unclear how the activation of Gi/o proteins affects the *I_h_* channel activity, and thus we only considered the Gs-activating effects of dopamine/norepinephrine and the M1-receptor-mediated (Gq-activating) effects of acetylcholine. Once more data is collected on how *I_h_* is modulated by different neuromodulatory receptors or intracellular cascades, the effects could be implemented following our procedure to model effects of acetylcholine and dopamine.

Our results showing that the *I_h_* channels facilitate APs for proximal stimulation and shunt distal stimulation raised important questions for the neuromodulation of L5PC. What are the effects of dopaminergic or cholinergic inputs at different parts of the dendritic tree on the excitability of the L5PCs? Can the effects of one neuromodulator at one part of a dendritic tree be compensated by another neuromodulator at other parts? Are there subclasses of L5PCs that respond differently to neuromodulators based on the distribution of their neuromodulatory receptors? Here, we further explored how these neuromodulatory actions on *I_h_* channels contribute separately or jointly to L5PC excitability.

When assessing the effect of dopaminergic modulation, our results showed that in the presence of a hot zone of Ca^2+^ channels, dopamine modulation decreased L5PC excitability when stimulated at the distal apical dendrite. This is consistent with the proposed role of D1-receptor-mediated effects of dopamine in the prefrontal cortex. D1 agonists increase HCN channels’ open probability. During normal brain processing state, this effect would help reduce the influence of lateral input to superficial layers in order to facilitate the processing of the neuron’s preferred stimulus [Avery and Krichmar, 2017] and enhance sustained firing during working memory tasks [Arnsten, 2011, Gamo et al., 2015].

A study on the effect of acetylcholine on dendro-somal integration in L5PCs found that optogenetic stimulus-evoked release of acetylcholine led to specific modulation of the apical dendrite [Williams and Fletcher, 2019]. In particular, when paired to both somatic and dendritic stimulation, it was found to greatly augment its effect in the somatic excitability. Here, we were able to explain this differential modulation: the presence of a hot zone of Ca^2+^ channels determines how the site of incoming stimulation will interact with *I_h_* currents.

Our computational modelling framework provided an efficient tool for flexibly studying the effects of different scenarios with combined neuromodulation. We collected our model predictions for L5PCs in Table 2. Our simulations with a combination of dopaminergic neuromodulation of *I_h_* channels in the distal apical dendrite and cholinergic neuromodulation in the proximal dendrite yielded stronger shunting of distal apical inputs than either of these modulations alone (and likewise, acetylcholine in the distal apical dendrite and dopamine in the proximal dendrite facilitated the neuron response to distal apical stimuli than either neuromodulator alone; Fig. 6). Our results obtained with the two L5PCs models strongly suggest that the *I_h_*-mediated shunting of distal inputs requires the hot zone of LVA Ca^2+^ channels: without the hot zone, the *I_h_* neuromodulation of the proximal apical dendrite determines whether the L5PC excitability is increased (dopamine) or decreased (acetylcholine), and the opposite neuromodulation at the distal apical dendrite only has a mildly compensating effect on the L5PC excitability (Fig. S6B,E). Moreover, our simulations suggest that the apical dendritic regime where dopaminergic modulation of *I_h_* decreases the L5PC excitability is narrowed down by simultaneous glutamatergic stimulation and expanded by simultaneous GABAergic inhibition of the basal dendrites (Fig. 5).

**Table 2:**
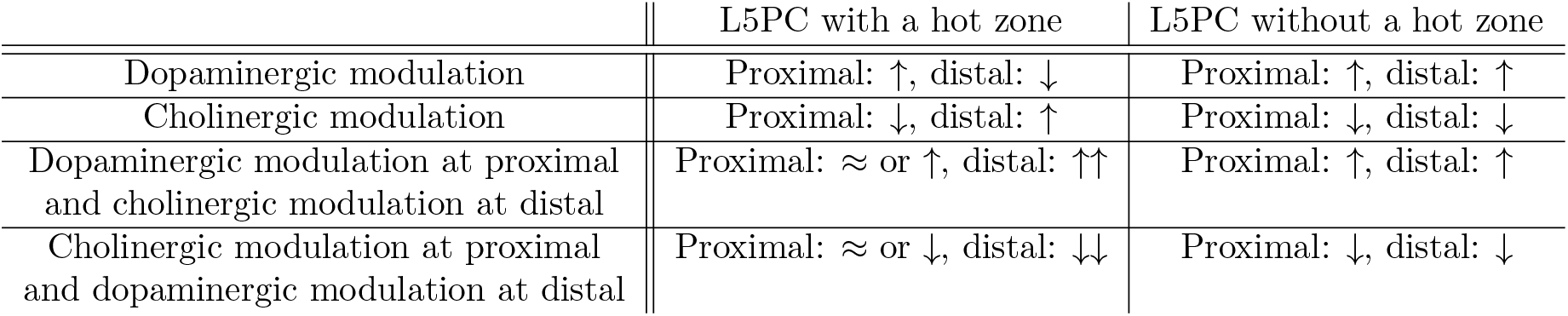
Predictions for how the excitability of L5PCs in response to proximal and distal apical dendrites is changed by modulation of *I_h_* channels. Higher excitability (decreased AP threshold) is denoted by ↑, and lower excitability (increased AP threshold) is denoted by ↓. The combinations ⇈ and ⇊ represent cases where the combination of cholinergic and dopaminergic modulations at different parts of the dendrite caused stronger increase or decrease to the L5PC excitability than any of the two neuromodulations alone.

Our results have functional implications, because the proximal and distal parts of the apical dendrite receive neuromodulatory inputs from different sources [Bloem et al., 2014, Aransay et al., 2015]. In addition, although experimental approaches often study the effect of single neuromodulatory systems, our findings are in line with the view that the combination of several systems, which is the most likely scenario during normal brain processing, may exert synergistic effects on neuron excitability [Levin et al., 1990]. Our work makes very specific predictions about the interaction between neuromodulators, input region and *I_h_* modulation of neuron excitability, which will be further extended once more data is gathered on the activity in the neuromodulatory sources and their interaction with each other and the cortex.

### 4.2 Comparison with previous computational studies of *I_h_* function on pyramidal cells

Of the previous computational studies, the approach of [Kase and Imoto, 2012] was methodologically closest to ours although they used a simplified, generic neuron model. Similar to our work, [Kase and Imoto, 2012] did not assume a coupling of *I_h_* with another current, yet they showed that the interactions between *I_h_* and two other channels, namely M-type K^+^ channels and T-type Ca^2+^ channels may crucially change the effects of blockage of *I_h_* on neuron excitability. However, they only analyzed these interactions for responses to somatic stimuli, while here we studied the responses to dendritic stimuli at different locations. In particular, they showed that by increasing the M-type K^+^ channel conductance the effect of *I_h_* blockade was changed from loss to gain of excitability [Kase and Imoto, 2012]. In their simulations, the change of T-type Ca^2+^ channel conductance did not show similar switch — however, this is most likely due to the high M-type K^+^ channel conductance or small range of T-type Ca^2+^ channel conductance chosen for these experiments, since the derivatives of the threshold currents with respect to the T-type Ca^2+^ channel conductance of *I_h_*-blocked and unblocked neurons are visibly different (Fig. 5f of [Kase and Imoto, 2012]). Our models, by contrast, predicted that M-type K^+^ channel conductance had little effect on the threshold currents both in absence and presence of *I_h_* currents (Fig. S2). This question begs for additional research, in particular when studying the effects of cholinergic neuromodulation of *I_h_*, since the M-type K^+^ channels are strongly modulated by acetylcholine [Brown and Adams, 1980]. Our models, however, highlight the crucial role of LVA (T-type) Ca^2+^ channels in whether *I_h_* channels promote or hinder L5PC excitability.

In another study on the role of *I_h_* current in CA1 py neurons, [Migliore et al., 2004] predicted that *I_h_* channels shunt synchronized but not unsynchronized inputs to CA1 pyramidal neurons. [George et al., 2009] suggested that the shunting effect seen in experiments of CA1 pyramidal neurons is caused by an interaction of *I_h_* and M-type K^+^ channels. However, [Migliore and Migliore, 2012] claimed that the conductance of the M-type K^+^ channels needed for shunting inhibition would be unrealistically high. Instead, they suggested that *I_h_* current is always coupled to another ionic current (i.e., when *I_h_* is blocked, the other current is blocked too) that drives the shunt-inhibition effects observed in experiments. [Kelley et al., 2021] examined a range of neocortical L5PC models, and suggested that a coupling of TASK-like channel with the *I_h_* currents as delineated in [Migliore and Migliore, 2012] provided the best fit to experimental impedance amplitude and phase data. While it is possible that different mechanisms modulate *I_h_* influence on neuron excitability on different regions, our models suggest that when using a realistically morphological model of an L5PC, T-type Ca^2+^ channels and not M-type K^+^ channels have a role in the shunting or excitatory effects of *I_h_* currents.

### 4.3 Relevance for cognition and behavior

Increasing evidence proposes the ability to integrate information within L5PCs as the mechanism behind some of the most complex processing capabilities of the brain. The L5PCs are unique in that inputs form sources with different information content are spatially segregated (for a review, see [Phillips et al., 2016]). Inputs to the basal dendrite or perisomatic zones are related to the preferred stimulus of the neuron, that is, the information itself that the neuron transmits as part of a feedforward circuit. On the other hand, inputs to the apical dendrite arrive from higher-order thalamus, feedback loops from prefrontal cortex, subcortical structures such as amygdala, and neuromodulatory regions [Gur and Snodderly, 2008, Rubio-Garrido et al., 2009]. These apical inputs modulate the strength of the effect that the input to the somatic compartment has on the neuron’s excitability. Thus, the apical inputs act as the context that exerts a modulation over the content that arrives to the basal input. For example, high arousal-related neuromodulatory activity enhances the transmission of salient stimuli in the neurons that process these specific stimuli, and thus provide the context that enhances the processing of important information. The ability to integrate context and content is at the core of higher-order brain functions. Brain activation during tasks and behavior is highly dependent on e.g. task goals, emotional or arousal state, and internal states determined by the autonomic system [Lee and Dan, 2012, Zagha and McCormick, 2014]. Our work adds to the existing literature on how this integration is implemented at the level of single neurons.

Recent theoretical and empirical work suggestss the mechanism of “apical amplification” [Marvan et al., 2021] or “dendritic integration” [Aru et al., 2020b] to be at the core of conscious processing. According to these views, only stimuli that are prioritized by contextual information reach the status of being consciously perceived. Furthermore, this cellular mechanism recapitulates the two main features of consciousness, meaning its level and content [Aru et al., 2019]. Arousal-related neuromodulatory systems like the ones studied here (i.e. noradrenaline, dopamine or acetylcholine) would set the level of consciousness by providing an alertness signal [Phillips et al., 2016]. The arousing or prioritizing effect of neuromodulatory signals (e.g. acetylcholine) is related to downregulation of *I_h_*, and was modelled here as a shift in the voltage dependence of *I_h_* current. Consistent with the apical amplification framework, we found an increase in neuron excitability in response to distal apical stimulation following this neuromodulatory effect. However, our simulations with combinations of basal and apical dendritic stimuli predicted that the more there is excitatory drive at the basal dendrite, the smaller the region in the apical dendrite where the *I_h_* current has the shunting effect (Fig. 5). Our results thus suggest that the *I_h_* current would more likely act as a shunt inhibitor in the mode of “apical drive” [Aru et al., 2020a] where the main excitatory drive arrives at the apical dendrite than in the mode of “apical amplification” where both apical and basal dendritic stimulation is needed for an AP.

Our ability to model the effects of multiple neuromodulatory systems on L5PC ability to integrate apical and basal inputs has special relevance in the mental conditions where consciousness is altered. For example, psychotic hallucinations, anesthesia, or dreaming are particular states where the integration of content is disconnected from the context [Phillips et al., 2016, Phillips et al., 2018, Aru et al., 2020a]. Such conditions are complex and challenging to describe, because what is altered seems to be the meaning or interpretation of the sensory processing, thus speaking of an impairment in the capacity to contextualize the information. Of the most common mental disorders with hallucinations, altered integration of context-dependent and sensory inputs to L5PCs could be a common pathophysiological feature particularly in schizophrenia, as suggested previously [Black et al., 2004, Arion et al., 2015, Mäki-Marttunen et al., 2016, Maki-Marttunen et al., 2018, Maki-Marttunen et al., 2019]. Our findings of the need of both LVA Ca^2+^ channels and *I_h_* currents for the shunting effect of the *I_h_* channels are interesting for schizophrenia research since both *I_h_* and LVA Ca^2+^ channels, alongside cholinergic and dopaminergic receptors, are products of risk genes of the mental disorder [Devor et al., 2017, Refisch et al., 2021, El Khoueiry et al., 2022]. Furthermore, neuromodulatory systems and their interaction likely play a crucial role in the pathophysiology of schizophrenia [Mäki-Marttunen et al., 2020]. Computational work like the one presented here offers a methodological approach to unify the different levels at which this and other mental disorders express their symptoms and phenotypes.

### 4.4 Future directions

In this work, we restricted our analysis on the contributions of *I_h_* currents to single-L5PC excitability. Previous computational modelling studies analyzed the effects of *I_h_* currents on network phenomena such as resonance to oscillations of different frequencies [Neymotin et al., 2013, Vaidya and Johnston, 2013]. *I_h_* current of L5PCs, and particularly the inactivation dynamics thereof, were also found important for local field potentials (LFP) in [Ness et al., 2016]. Our approach of studying the effects of *I_h_* neuromodulation could be directly applied to the analysis of network phenomena as well. In particular, the dependence of working memory-like network activity (as modelled in, e.g., [Compte et al., 2000]) on neuromodulation of *I_h_* channels could provide important insights into the mechanisms in which neuromodulators shape cognitive processes. Future models of *I_h_* activity could also benefit from integrating more biochemical interactions between the channels, in particular Ca^2+^ signalling, as done in [Neymotin et al., 2016].

## Acknowledgements

Funding: Academy of Finland (336376), Research Council of Norway (248828), and University of Oslo Convergence Environment (4MENT). UNINETT Sigma2 resources (project NN9529K) and CSC (project 2003397) were used for simulations.

## Supplementary figures and tables

**Figure S1:**
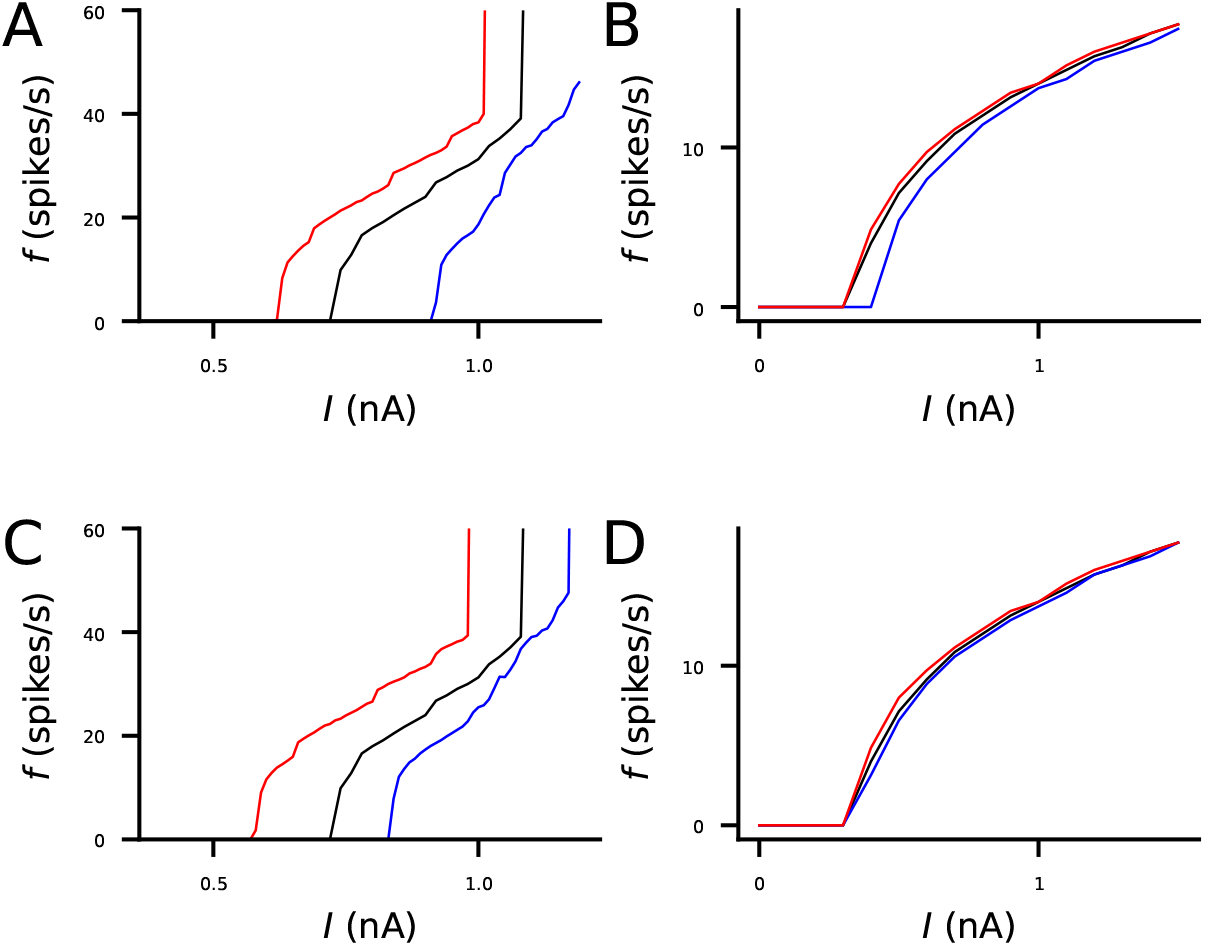
*I_h_* activation in the dendrites increases the frequency of action potentials in response to somatic DC in L5PCs. **A–B**: The frequency of APs (y-axis) in response to somatic DC of a given amplitude (x-axis) in Almog (A) and Hay (B) model neurons under up- or down-regulated *I_h_* channels. Black: control neuron. Blue: *I_h_* conductance blocked in the apical dendrite. Red: *I_h_* conductance increased by 100% in the apical dendrite. **C–D**: The frequency of APs response to somatic DC in Almog (C) and Hay (D) model neurons under different neuromodulatory states. Black: control neuron. Blue: cholinergic neuromodulation. Red: dopaminergic neuromodulation.

**Figure S2:**
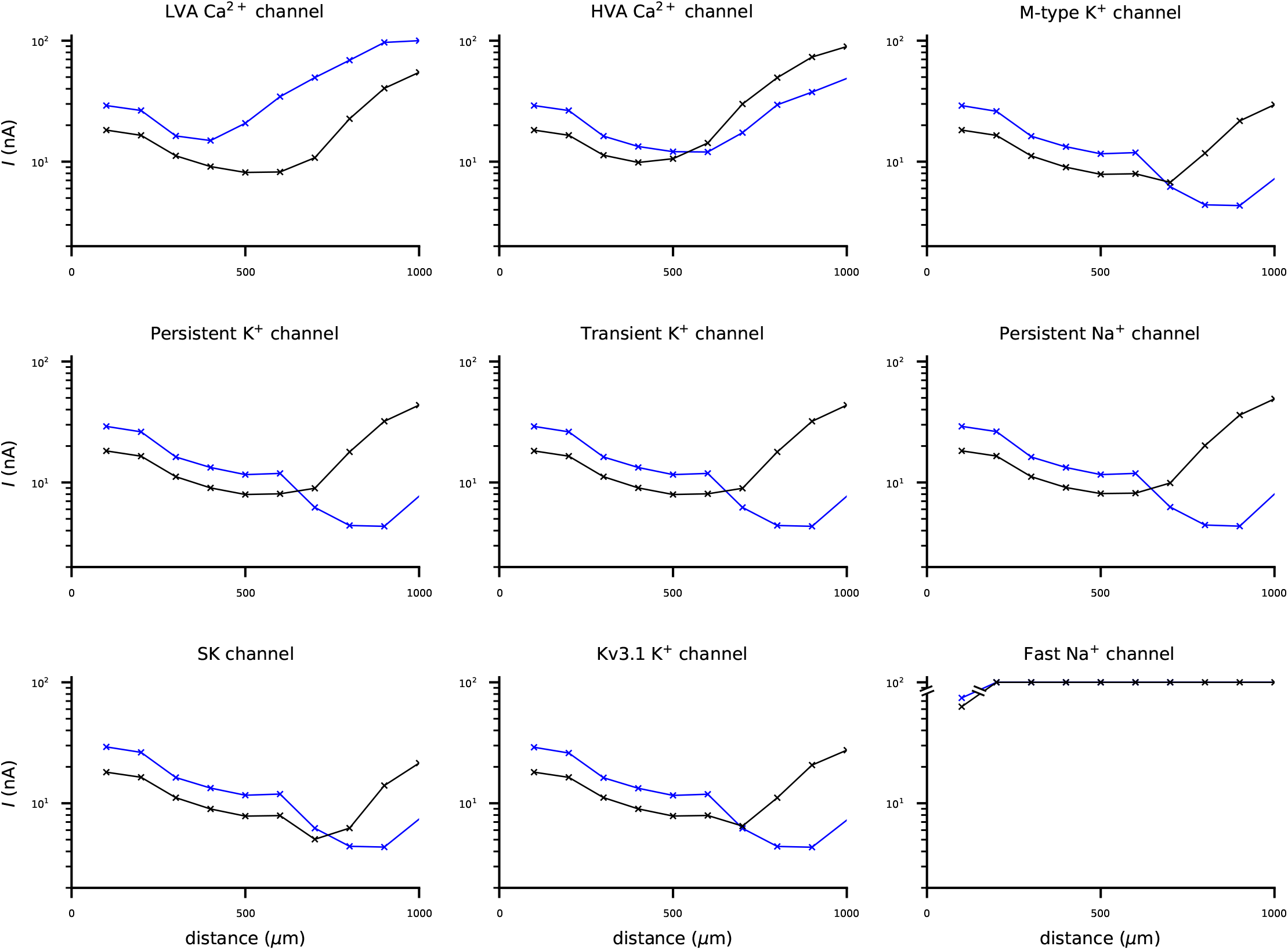
Blockage of fast Na^+^ channels abolishes spiking and blockage of LVA Ca^2+^ channels from the Hay model prohibits the shunting of distal apical dendritic stimuli by *I_h_* currents, but blockage of other ion channels does not affect the qualitative behaviour where *I_h_* currents make the neuron more excitable by strong proximal inputs and less excitable by strong distal inputs at the apical dendrite. See Fig. 2 for details. **A**: LVA Ca^2+^ channels blocked. **B**: HVA Ca^2+^ channels blocked. **C**: M-type K^+^ channels blocked. **D**: Persistent K^+^ channels blocked. **E**: Transient K^+^ channels blocked. **F**: Persistent Na^+^ channels blocked. **G**: Ca^2+^-dependent K^+^ channels (SK channels) blocked. **H**: Kv3.1-type K^+^ channels blocked. **F**: Transient Na^+^ channels blocked. Black curves: the named ion channel blocked. Blue curves: the named ion channel and the *I_h_* channel blocked.

**Figure S3:**
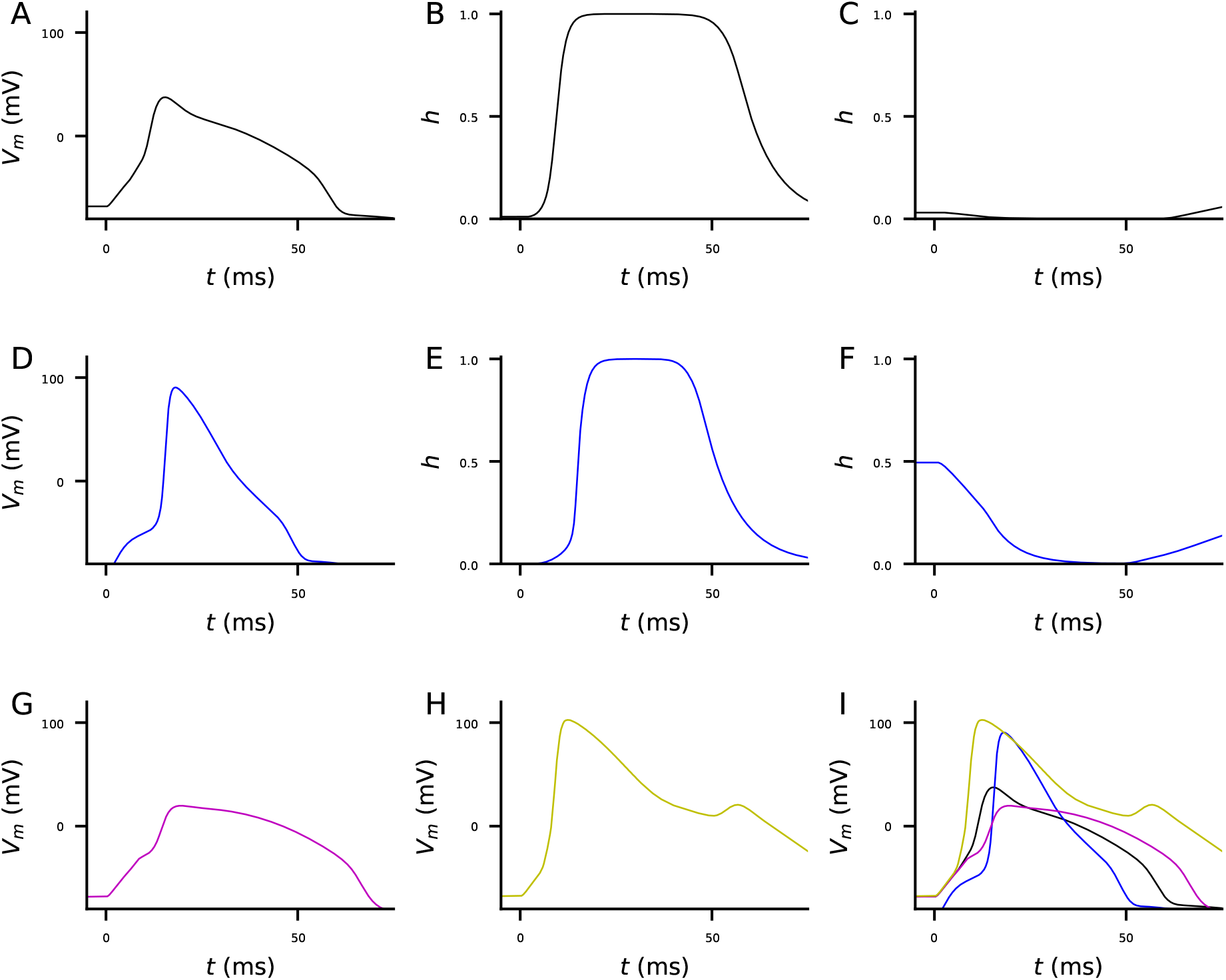
A single-compartment model of a distal apical dendritic section of the Hay-model L5PC predicts that the shunting effect of *I_h_* currents is mediated by a high degree of LVA Ca^2+^ channel inactivation in the resting state in presence of *I_h_* currents.**A–C**: Time courses of the membrane potential (A) and the activation (B) and inactivation (C) variables of the LVA Ca^2+^ channels in the control condition when the compartment was stimulated with an alpha-shaped synaptic conductance of 2 nS. **D–F**: Time courses of the membrane potential (D) and the activation (E) and inactivation (G) variables when *I_h_* channels were blocked. **G**: Time course of the membrane potential of a model compartment, where the LVA Ca^2+^ current is replaced by an artificial LVA current species where the values of the activation variable are directly taken from *I_h_*-blocked simulation (E) and the those of the inactivation variable are taken from the control simulation (C). This model compartment produces a **milder** response than either the control (A) or *I_h_*-blocked neuron (D). **H**: Time course of the membrane potential of a model compartment, where the LVA Ca^2+^ current is replaced by an artificial LVA current species where activation variable is taken from the control simulation (B) and the inactivation variable is taken from the *I_h_*-blocked simulation (F). This model compartment produces a **stronger** response than either the control (A) or *I_h_*-blocked neuron (D). **I**: The membrane potential time courses from panels (A), (D), and (G-H) overlaid.

**Figure S4:**
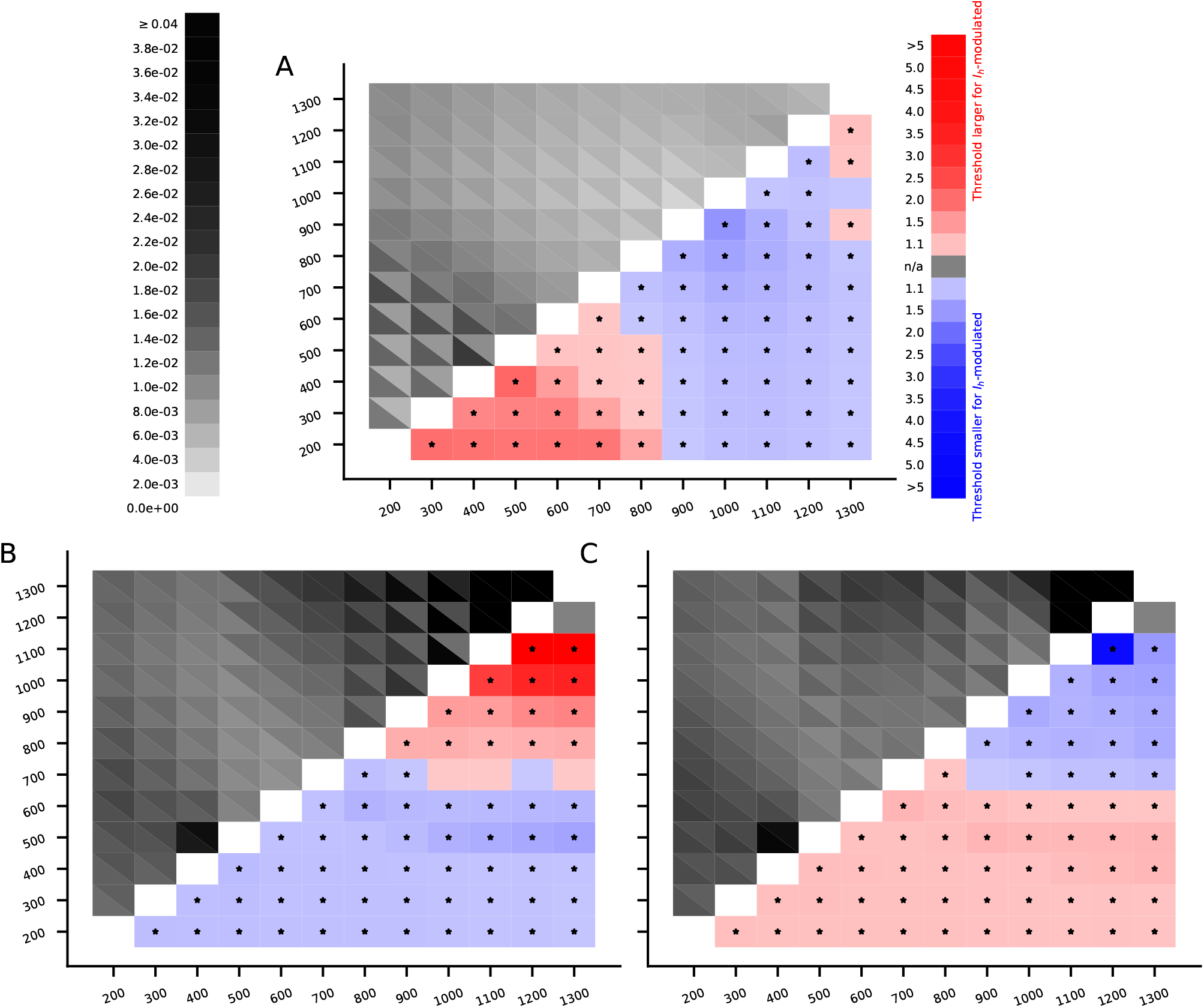
*I_h_* current-mediated shunting of distal apical dendritic stimuli shown by different simulations. **A**: Predictions of the Almog model with a hot zone of Ca^2+^ channels. The upper left grid shows the threshold conductances for a set of 2000 excitatory synapses to induce an AP, and the lower right grid shows the factor by which the threshold conductance of the *I_h_*-blocked neuron is larger (red) or smaller (blue) than that of the control neuron. See Fig. 4F for details. **B–C**: Predictions of the Hay model for the dopaminergically (B) or cholinergically (C) modulated compared to the non-modulated neuron. *Upper left grid*: The threshold conductances for a set of 2000 excitatory synapses to induce an AP in the Hay model. In each grid slot, the color of the upper right triangle indicates the threshold conductance in the control neuron wherease that of the lower left triangle indicates the threshold conductance in the dopaminergically (B) or cholinergically (C) modulated neuron. *Lower right grid*: The factor by which the threshold conductance of the dopaminergically (B) or cholinergically (C) modulated neuron is larger (red) or smaller (blue) than that of the control neuron.

**Figure S5:**
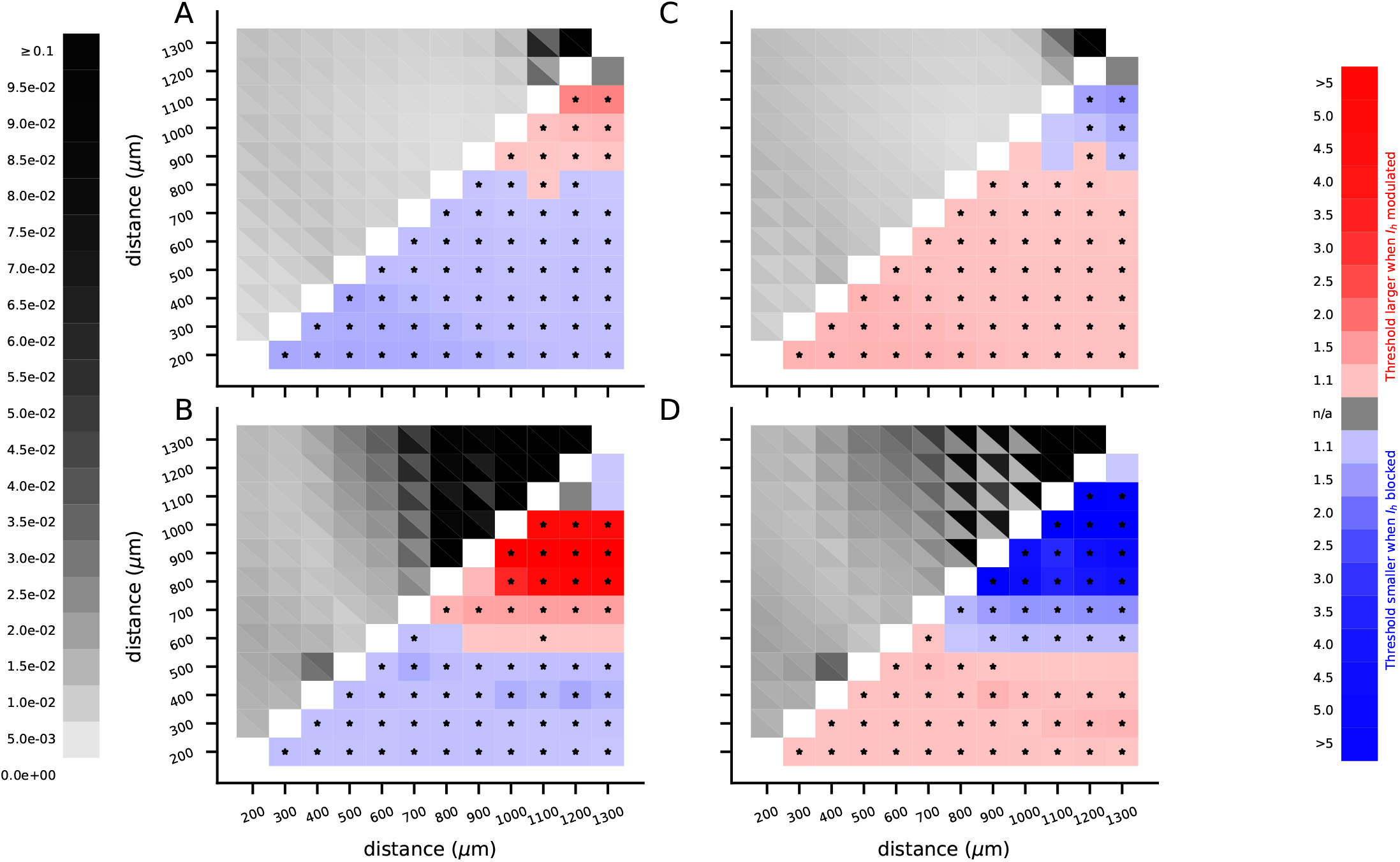
The shunting effect of dopaminergic neuromodulation and the excitability-increasing effect of cholinergic neuromodulation are further constrained to the distal apical dendrite by glutamatergic stimulation of the basal dendrite and expanded by GABAergic stimulation of the basal dendrite. The four panels show the Hay-model predictions for the effects of dopaminergic (A,C) or cholinergic (B,D) neuromodulation on the AP thresholds of apical dendritic stimulation in the presence of simultaneous glutamatergic (A–B) or GABAergic (C–D) stimulation of the basal dendrite. *Upper left grids*: The threshold conductances for a set of 2000 excitatory synapses to induce an AP in the Hay model. In each grid slot, the color of the upper right triangle indicates the threshold conductance in the control neuron wherease that of the lower left triangle indicates the threshold conductance in the dopaminergically (A–B) or cholinergically (C–D) modulated neuron. *Lower right grids*: The factor by which the threshold conductance of the dopaminergically (A,C) or cholinergically (B,D) modulated neuron is larger (red) or smaller (blue) than that of the control neuron.

**Figure S6:**
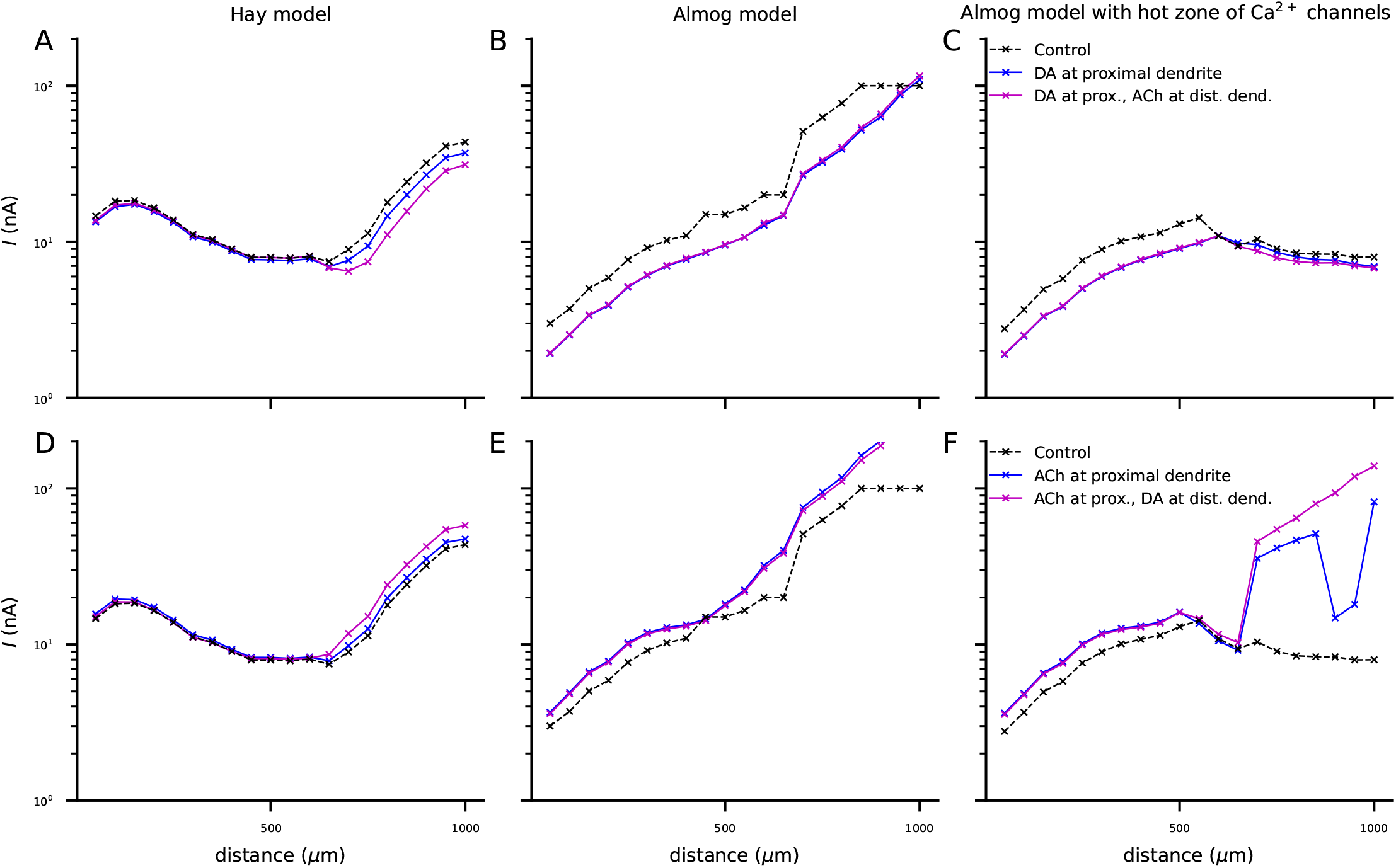
Combination of dopaminergic and cholinergic neuromodulation can increase or decrease the AP threshold throughout the apical dendrite in L5PCs expressing a hot zone of LVA Ca^2+^ channels. **A–C**: Threshold amplitude for a 2-ms current input applied to the apical dendrite at a given distance (x-axis) from soma according to Hay model (A), Almog model (B), or Almog model with a hot zone of LVA Ca^2+^ channels (C). Black: control neuron. Blue: neuron with proximal apical dendrite modulated by dopamine. Magenta: neuron with proximal apical dendrite modulated by dopamine and distal apical dendrite by acetylcholine. **D–F**: The experiment of (A)–(C) repeated with opposite modulation, i.e., control neuron (black) and neuron with cholinergic modulation of proximal apical dendrite with (magenta) or without (blue) dopaminergic modulation of the distal apical dendrite.

## References

Aćimović, J., Mäki-Marttunen, T., Teppola, H., and Linne, M.-L. (2021). Analysis of cellular and synaptic mechanisms behind spontaneous cortical activity in vitro: Insights from optimization of spiking neuronal network models. bioRxiv, page 466340.

Almog, M. and Korngreen, A. (2014). A quantitative description of dendritic conductances and its application to dendritic excitation in layer 5 pyramidal neurons. Journal of Neuroscience, 34(1):182–196.

Aransay, A., Rodríguez-López, C., García-Amado, M., Clascá, F., and Prensa, L. (2015). Long-range projection neurons of the mouse ventral tegmental area: a single-cell axon tracing analysis. Frontiers in neuroanatomy, 9:59.

Arion, D., Corradi, J. P., Tang, S., Datta, D., Boothe, F., He, A., Cacace, A. M., Zaczek, R., Albright, C. F., Tseng, G., et al. (2015). Distinctive transcriptome alterations of prefrontal pyramidal neurons in schizophrenia and schizoaffective disorder. Molecular psychiatry, 20(11):1397–1405.

Arnsten, A. F. (2011). Catecholamine influences on dorsolateral prefrontal cortical networks. Biological psychiatry, 69(12):e89–e99.

Aru, J., Siclari, F., Phillips, W. A., and Storm, J. F. (2020a). Apical drive-a cellular mechanism of dreaming? Neuroscience & Biobehavioral Reviews, 119:440–455.

Aru, J., Suzuki, M., and Larkum, M. E. (2020b). Cellular mechanisms of conscious processing. Trends in Cognitive Sciences, 24(10):814–825.

Aru, J., Suzuki, M., Rutiku, R., Larkum, M. E., and Bachmann, T. (2019). Coupling the state and contents of consciousness. Frontiers in Systems Neuroscience, page 43.

Avery, M. C. and Krichmar, J. L. (2017). Neuromodulatory systems and their interactions: a review of models, theories, and experiments. Frontiers in neural circuits, 11:108.

Ballo, A. W., Keene, J. C., Troy, P. J., Goeritz, M. L., Nadim, F., and Bucher, D. (2010). Dopamine modulates ih in a motor axon. Journal of Neuroscience, 30(25):8425–8434.

Berger, T., Larkum, M. E., and Lüscher, H.-R. (2001). High ih channel density in the distal apical dendrite of layer v pyramidal cells increases bidirectional attenuation of epsps. Journal of neurophysiology, 85(2):855–868.

Bergson, C., Mrzljak, L., Smiley, J. F., Pappy, M., Levenson, R., and Goldman-Rakic, P. S. (1995). Regional, cellular, and subcellular variations in the distribution of d1 and d5 dopamine receptors in primate brain. Journal of Neuroscience, 15(12):7821–7836.

Black, J. E., Kodish, I. M., Grossman, A. W., Klintsova, A. Y., Orlovskaya, D., Vostrikov, V., Uranova, N., and Greenough, W. T. (2004). Pathology of layer v pyramidal neurons in the prefrontal cortex of patients with schizophrenia. American Journal of Psychiatry, 161(4):742–744.

Bloem, B., Schoppink, L., Rotaru, D. C., Faiz, A., Hendriks, P., Mansvelder, H. D., van de Berg, W. D., and Wouterlood, F. G. (2014). Topographic mapping between basal forebrain cholinergic neurons and the medial prefrontal cortex in mice. Journal of Neuroscience, 34(49):16234–16246.

Branchereau, P., Van Bockstaele, E. J., Chan, J., and Pickel, V. M. (1996). Pyramidal neurons in rat prefrontal cortex show a complex synaptic response to single electrical stimulation of the locus coeruleus region: evidence for antidromic activation and gabaergic inhibition using in vivo intracellular recording and electron microscopy. Synapse, 22(4):313–331.

Brown, D. and Adams, P. (1980). Muscarinic suppression of a novel voltage-sensitive k+ current in a vertebrate neurone. Nature, 283(5748):673–676.

Byczkowicz, N., Eshra, A., Montanaro, J., Trevisiol, A., Hirrlinger, J., Kole, M. H., Shigemoto, R., and Hallermann, S. (2019). Hcn channel-mediated neuromodulation can control action potential velocity and fidelity in central axons. Elife, 8:e42766.

Compte, A., Brunel, N., Goldman-Rakic, P. S., and Wang, X.-J. (2000). Synaptic mechanisms and network dynamics underlying spatial working memory in a cortical network model. Cerebral cortex, 10(9):910–923.

Devor, A., Andreassen, O. A., Wang, Y., Mäki-Marttunen, T., Smeland, O. B., Fan, C.-C., Schork, A. J., Holland, D., Thompson, W. K., Witoelar, A., Chen, C.-C., Desikan, R. S., McEvoy, L. K., Djurovic, S., Greengard, P., Svenningsson, P., Einevoll, G. T., and Dale, A. M. (2017). Genetic evidence for role of integration of fast and slow neurotransmission in schizophrenia. Molecular Psychiatry (in press).

El Khoueiry, C., Cabungcal, J.-H., Rovó, Z., Fournier, M., Do, K. Q., and Steullet, P. (2022). Developmental oxidative stress leads to t-type ca2+ channel hypofunction in thalamic reticular nucleus of mouse models pertinent to schizophrenia. Molecular Psychiatry, pages 1–10.

Endo, T., Tarusawa, E., Notomi, T., Kaneda, K., Hirabayashi, M., Shigemoto, R., and Isa, T. (2008). Dendritic i h ensures high-fidelity dendritic spike responses of motion-sensitive neurons in rat superior colliculus. Journal of neurophysiology, 99(5):2066–2076.

Gamo, N. J., Lur, G., Higley, M. J., Wang, M., Paspalas, C. D., Vijayraghavan, S., Yang, Y., Ramos, B. P., Peng, K., Kata, A., et al. (2015). Stress impairs prefrontal cortical function via d1 dopamine receptor interactions with hyperpolarization-activated cyclic nucleotide-gated channels. Biological psychiatry, 78(12):860–870.

Gee, S., Ellwood, I., Patel, T., Luongo, F., Deisseroth, K., and Sohal, V. S. (2012). Synaptic activity unmasks dopamine d2 receptor modulation of a specific class of layer v pyramidal neurons in prefrontal cortex. Journal of Neuroscience, 32(14):4959–4971.

George, M. S., Abbott, L., and Siegelbaum, S. A. (2009). Hcn hyperpolarization-activated cation channels inhibit epsps by interactions with m-type k+ channels. Nature neuroscience, 12(5):577–584.

Gulledge, A. T. and Jaffe, D. B. (1998). Dopamine decreases the excitability of layer v pyramidal cells in the rat prefrontal cortex. Journal of Neuroscience, 18(21):9139–9151.

Gur, M. and Snodderly, D. M. (2008). Physiological differences between neurons in layer 2 and layer 3 of primary visual cortex (v1) of alert macaque monkeys. The Journal of physiology, 586(9):2293–2306.

Hay, E., Hill, S., Schürmann, F., Markram, H., and Segev, I. (2011). Models of neocortical layer 5b pyramidal cells capturing a wide range of dendritic and perisomatic active properties. PLoS Computational Biology, 7:e1002107.

He, C., Chen, F., Li, B., and Hu, Z. (2014). Neurophysiology of hcn channels: from cellular functions to multiple regulations. Progress in neurobiology, 112:1–23.

Hestrin, S. (1987). The properties and function of inward rectification in rod photoreceptors of the tiger salamander. The Journal of physiology, 390(1):319–333.

Kase, D. and Imoto, K. (2012). The role of hcn channels on membrane excitability in the nervous system. Journal of signal transduction, 2012.

Kelley, C., Dura-Bernal, S., Neymotin, S. A., Antic, S. D., Carnevale, N. T., Migliore, M., and Lytton, W. W. (2021). Effects of ih and task-like shunting current on dendritic impedance in layer 5 pyramidal-tract neurons. Journal of Neurophysiology, 125(4):1501–1516.

Labarrera, C., Deitcher, Y., Dudai, A., Weiner, B., Amichai, A. K., Zylbermann, N., and London, M. (2018). Adrenergic modulation regulates the dendritic excitability of layer 5 pyramidal neurons in vivo. Cell reports, 23(4):1034–1044.

Larkum, M. (2013). A cellular mechanism for cortical associations: An organizing principle for the cerebral cortex. Trends in Neurosciences, 36(3):141–151.

Lee, S.-H. and Dan, Y. (2012). Neuromodulation of brain states. Neuron, 76(1):209–222.

Levin, E. D., McGurk, S. R., Rose, J. E., and Butcher, L. L. (1990). Cholinergic-dopaminergic interactions in cognitive performance. Behavioral and neural biology, 54(3):271–299.

Maccaferri, G. and McBain, C. J. (1996). The hyperpolarization-activated current (ih) and its contribution to pacemaker activity in rat ca1 hippocampal stratum oriens-alveus interneurones. The Journal of physiology, 497(1):119–130.

Magee, J. C. (1999). Dendritic i h normalizes temporal summation in hippocampal ca1 neurons. Nature neuroscience, 2(6):508–514.

Mäki-Marttunen, T., Devor, A., Phillips, W. A., Dale, A. M., Andreassen, O. A., and Einevoll, G. T. (2019). Computational modeling of genetic contributions to excitability and neural coding in layer v pyramidal cells: applications to schizophrenia pathology. Frontiers in computational neuroscience, 13:66.

Mäki-Marttunen, T., Halnes, G., Devor, A., Witoelar, A., Bettella, F., Djurovic, S., Wang, Y., Einevoll, G. T., Andreassen, O. A., and Dale, A. M. (2016). Functional effects of schizophrenia-linked genetic variants on intrinsic single-neuron excitability: A modeling study. Biological Psychiatry: Cognitive Neuroscience and Neuroimaging, 1.

Mäki-Marttunen, T., Krull, F., Bettella, F., Hagen, E., Næss, S., Ness, T. V., Moberget, T., Elvsåshagen, T., Metzner, C., Devor, A., Edwards, A. G., Fyhn, M., Djurovic, S., Dale, A. M., Andreassen, O. A., and Einevoll, G. T. (2018). Alterations in schizophrenia-associated genes can lead to increased power in delta oscillations. Cerebral Cortex, 29(2):875–891.

Mäki-Marttunen, T., Lines, G. T., Edwards, A. G., Tveito, A., Dale, A. M., Einevoll, G. T., and Andreassen, O. A. (2017). Pleiotropic effects of schizophrenia-associated genetic variants in neuron firing and cardiac pacemaking revealed by computational modeling. Translational Psychiatry, 7(11):5.

Mäki-Marttunen, V., Andreassen, O. A., and Espeseth, T. (2020). The role of norepinephrine in the pathophysiology of schizophrenia. Neuroscience & Biobehavioral Reviews, 118:298–314.

Marvan, T., Polák, M., Bachmann, T., and Phillips, W. A. (2021). Apical amplification-a cellular mechanism of conscious perception? Neuroscience of consciousness, 2021(2):niab036.

McCormick, D. A. and Pape, H.-C. (1990). Properties of a hyperpolarization-activated cation current and its role in rhythmic oscillation in thalamic relay neurones. The Journal of physiology, 431(1):291–318.

Migliore, M., Messineo, L., and Ferrante, M. (2004). Dendritic i h selectively blocks temporal summation of unsynchronized distal inputs in ca1 pyramidal neurons. Journal of computational neuroscience, 16(1):5–13.

Migliore, M. and Migliore, R. (2012). Know your current ih: interaction with a shunting current explains the puzzling effects of its pharmacological or pathological modulations. PloS one, 7(5):e36867.

Mrzljak, L., Levey, A. I., Belcher, S., and Goldman-Rakic, P. (1998). Localization of the m2 muscarinic acetylcholine receptor protein and mrna in cortical neurons of the normal and cholinergically deafferented rhesus monkey. Journal of Comparative Neurology, 390(1):112–132.

Mrzljak, L., Levey, A. I., and Goldman-Rakic, P. S. (1993). Association of m1 and m2 muscarinic receptor proteins with asymmetric synapses in the primate cerebral cortex: morphological evidence for cholinergic modulation of excitatory neurotransmission. Proceedings of the National Academy of Sciences, 90(11):5194–5198.

Ness, T. V., Remme, M. W., and Einevoll, G. T. (2016). Active subthreshold dendritic conductances shape the local field potential. The Journal of physiology, 594(13):3809–3825.

Neymotin, S. A., Hilscher, M. M., Moulin, T. C., Skolnick, Y., Lazarewicz, M. T., and Lytton, W. W. (2013). Ih tunes theta/gamma oscillations and cross-frequency coupling in an in silico ca3 model. PLoS One, 8(10):e76285.

Neymotin, S. A., McDougal, R. A., Bulanova, A. S., Zeki, M., Lakatos, P., Terman, D., Hines, M. L., and Lytton, W. W. (2016). Calcium regulation of hcn channels supports persistent activity in a multiscale model of neocortex. Neuroscience, 316:344–366.

Oda, S., Tsuneoka, Y., Yoshida, S., Adachi-Akahane, S., Ito, M., Kuroda, M., and Funato, H. (2018). Immunolocalization of muscarinic m1 receptor in the rat medial prefrontal cortex. Journal of Comparative Neurology, 526(8):1329–1350.

O’Donnell, J., Zeppenfeld, D., McConnell, E., Pena, S., and Nedergaard, M. (2012). Norepinephrine: a neuromodulator that boosts the function of multiple cell types to optimize cns performance. Neurochemical research, 37(11):2496–2512.

Pedarzani, P. and Storm, J. F. (1995). Protein kinase a-independent modulation of ion channels in the brain by cyclic amp. Proceedings of the National Academy of Sciences, 92(25):11716–11720.

Phillips, W., Larkum, M. E., Harley, C. W., and Silverstein, S. M. (2016). The effects of arousal on apical amplification and conscious state. Neuroscience of consciousness, 2016(1).

Phillips, W. A., Bachmann, T., and Storm, J. F. (2018). Apical function in neocortical pyramidal cells: a common pathway by which general anesthetics can affect mental state. Frontiers in neural circuits, 12:50.

Poolos, N. P., Migliore, M., and Johnston, D. (2002). Pharmacological upregulation of h-channels reduces the excitability of pyramidal neuron dendrites. Nature neuroscience, 5(8):767–774.

Radnikow, G. and Feldmeyer, D. (2018). Layer-and cell type-specific modulation of excitatory neuronal activity in the neocortex. Frontiers in neuroanatomy, 12:1.

Refisch, A., Chung, H.-Y., Komatsuzaki, S., Schumann, A., Mühleisen, T. W., Nöthen, M. M., Huebner, C. A., and Bär, K.-J. (2021). A common variation in hcn1 is associated with heart rate variability in schizophrenia. Schizophrenia research, 229:73–79.

Rubio-Garrido, P., Pérez-de Manzo, F., Porrero, C., Galazo, M. J., and Clascá, F. (2009). Thalamic input to distal apical dendrites in neocortical layer 1 is massive and highly convergent. Cerebral cortex, 19(10):2380–2395.

Shine, J. M., Breakspear, M., Bell, P. T., Ehgoetz Martens, K. A., Shine, R., Koyejo, O., Sporns, O., and Poldrack, R. A. (2019). Human cognition involves the dynamic integration of neural activity and neuromodulatory systems. Nature neuroscience, 22(2):289–296.

Shiozaki, K., Iseki, E., Hino, H., and Kosaka, K. (2001). Distribution of m1 muscarinic acetylcholine receptors in the hippocampus of patients with alzheimer’s disease and dementia with lewy bodies-an immunohistochemical study. Journal of the neurological sciences, 193(1):23–28.

Tsay, D., Dudman, J. T., and Siegelbaum, S. A. (2007). Hcn1 channels constrain synaptically evoked ca2+ spikes in distal dendrites of ca1 pyramidal neurons. Neuron, 56(6):1076–1089.

Turrini, P., Casu, M. A., Wong, T. P., De Koninck, Y., Ribeiro-da Silva, A., and Cuello, A. C. (2001). Cholinergic nerve terminals establish classical synapses in the rat cerebral cortex: synaptic pattern and age-related atrophy. Neuroscience, 105(2):277–285.

Vaidya, S. P. and Johnston, D. (2013). Temporal synchrony and gamma-to-theta power conversion in the dendrites of ca1 pyramidal neurons. Nature neuroscience, 16(12):1812–1820.

Vasquez, C. and Lewis, D. (2003). The β2-adrenergic receptor specifically sequesters gs but signals through both gs and gi/o in rat sympathetic neurons. Neuroscience, 118(3):603–610.

Williams, S. R. and Fletcher, L. N. (2019). A dendritic substrate for the cholinergic control of neocortical output neurons. Neuron, 101(3):486–499.

Xing, B., Li, Y.-C., and Gao, W.-J. (2016). Norepinephrine versus dopamine and their interaction in modulating synaptic function in the prefrontal cortex. Brain research, 1641:217–233.

Zagha, E. and McCormick, D. A. (2014). Neural control of brain state. Current opinion in neurobiology, 29:178–186.

Zhao, Z., Zhang, K., Liu, X., Yan, H., Ma, X., Zhang, S., Zheng, J., Wang, L., and Wei, X. (2016). Involvement of hcn channel in muscarinic inhibitory action on tonic firing of dorsolateral striatal cholinergic interneurons. Frontiers in cellular neuroscience, 10:71.

Zhou, H.-C., Sun, Y.-Y., Cai, W., He, X.-T., Yi, F., Li, B.-M., and Zhang, X.-H. (2013). Activation of *β*2-adrenoceptor enhances synaptic potentiation and behavioral memory via camp-pka signaling in the medial prefrontal cortex of rats. Learning & Memory, 20(5):274–284.

